# An Accelerometer Based Heart Monitor to Measure Changes of the Autonomic Nervous System

**DOI:** 10.1101/2022.06.02.494453

**Authors:** Viktoriya Babenko, Neil M. Dundon, Alan Macy, Alexandra Stump, Macey Turbow, Matthew Cieslak, Scott T. Grafton

## Abstract

The electrocardiogram (ECG) and impedance cardiography (ICG) are typically combined to estimate electromechanical features such as the pre-ejection period (PEP) and left ventricular ejection time (LVET); indicators of changes in the cardiac specific drive of the autonomic nervous system (ANS). Current methods of ICG are time intensive in subject preparation and the measurements are vulnerable to non-reproducible subject-specific electrode configuration. Furthermore, analysis of impedance waveforms can be time consuming and labeling of key time points can suffer from experimenter bias. Here we present a wearable heart monitor that includes ECG, but replaces the commonly used 8 ICG electrodes with a single accelerometer (ACC) placed at the suprasternal notch. The ACC indirectly measures movement of the arterial pulse wave as blood is ejected into the aorta and great vessels. The resulting ACC waveform is processed into two smooth and readily identified waves, corresponding to the timing of the opening and closing of the aortic valve. We tested the ACC’s utility and reliability for tracking cardiac ANS tone by comparing PEP and LVET measurements obtained simultaneously with conventional ICG and the ACC. Participants were recorded in the sitting and supine position with ECG, ICG, and ACC. While seated, they engaged in a classic physical stress task known to modulate ANS activity. There were obvious and significant associations between ICG and ACC estimates of PEP and LVET derivatives with respect to time. These findings support ACC as a complementary method for tracking ANS that is robust, time efficient, and readily accessible to researchers.

## 1 INTRODUCTION

Research quantifying autonomic nervous system activity (ANS) and cardiac mobilization is becoming increasingly important and prevalent - not only in medical domains, but also in psychological and brain sciences (Cybulski et al., 2012; Thayer et al., 2010). These measures provide insight into motivational states, stress reactivity, reward sensitivity, task engagement and decision making (Kuipers et al., 2017; Richter et al., 2008; Richter & Gendolla, 2007, 2009). There are various methods in use to measure the psychophysiological response of the ANS such as skeletomuscular activation, hormonal fluctuations, systolic blood pressure dynamics and assays of the immune response (Blascovich & Mendes, 2010). The most common noninvasive methods include measures of cardiovascular reactivity from electrocardiography (ECG) to estimate heart rate variability (HRV). High frequency HRV is associated with parasympathetic tone. While low frequency HRV is claimed to be an indicator of a sympathetic tone, recent work establishes that it is an unreliable index, due to influences by both sympathetic and parasympathetic activity (Berntson et al., 1997; Reyes del Paso et al., 2013; Valenza et al., 2018). As an alternative to HRV, when ECG is combined with impedance cardiography (ICG) it is possible to extract temporal indices of cardiac function sensitive to autonomic tone such as pre-ejection period (PEP) and left-ventricular ejection time (LVET).

PEP (Figure 1) is the sum of the electromechanical delay and isovolumic contraction time (contraction of the ventricle prior to opening of the aortic valve) (Tomaka et al., 1997; Kelsey et al., 2004; Wright & Kirby, 2001). PEP in particular has been shown to be a reliable indicator of the sympathetic nervous system (SNS), resulting in an inverse relationship where a decrease in PEP represents an increase in sympathetic activity. PEP is sensitive to the predictive effects of task manipulations (Kelsey et al., 2000, 2004), decision making (Dundon et al., 2020, 2021) and individual differences in a variety of tasks (Kelsey, 1991; Kelsey et al., 2001). LVET is defined by the time interval between the opening and closing of the aortic valve, representing the interval during which the left ventricle ejects blood into the aorta (Figure 1). LVET demonstrates a trend of decreasing duration with increased sympathetic tone. However, it also provides insight into the preload effects on the heart (Uijtdehaage & Thayer, 2000). As intrathoracic pressure is changed during baroreflex, the preload effects will vary with LVET (Hassan & Turner, 1983). This mechanism is influenced by the Frank-Starling effect (Spodick, 1979). Thus, unlike PEP, it is not considered a measure that isolates SNS activity. When continuous blood pressure (CBP) is available it is also possible to combine LVET with models of the chest to estimate cardiac output (CO) and total peripheral vascular resistance (Kelsey et al., 2004; Kreibig et al., 2013; Matthews et al., 2003; Seery, 2013; Wright et al., 1986). These model based measures have been used broadly to study baseline levels of health (Kelsey, 2004) and to test the biopsychosocial model of challenge and threat to distinguish between varying states of motivation (Blascovich, 2008; Tomaka et al., 1997).

**FIGURE 1.**
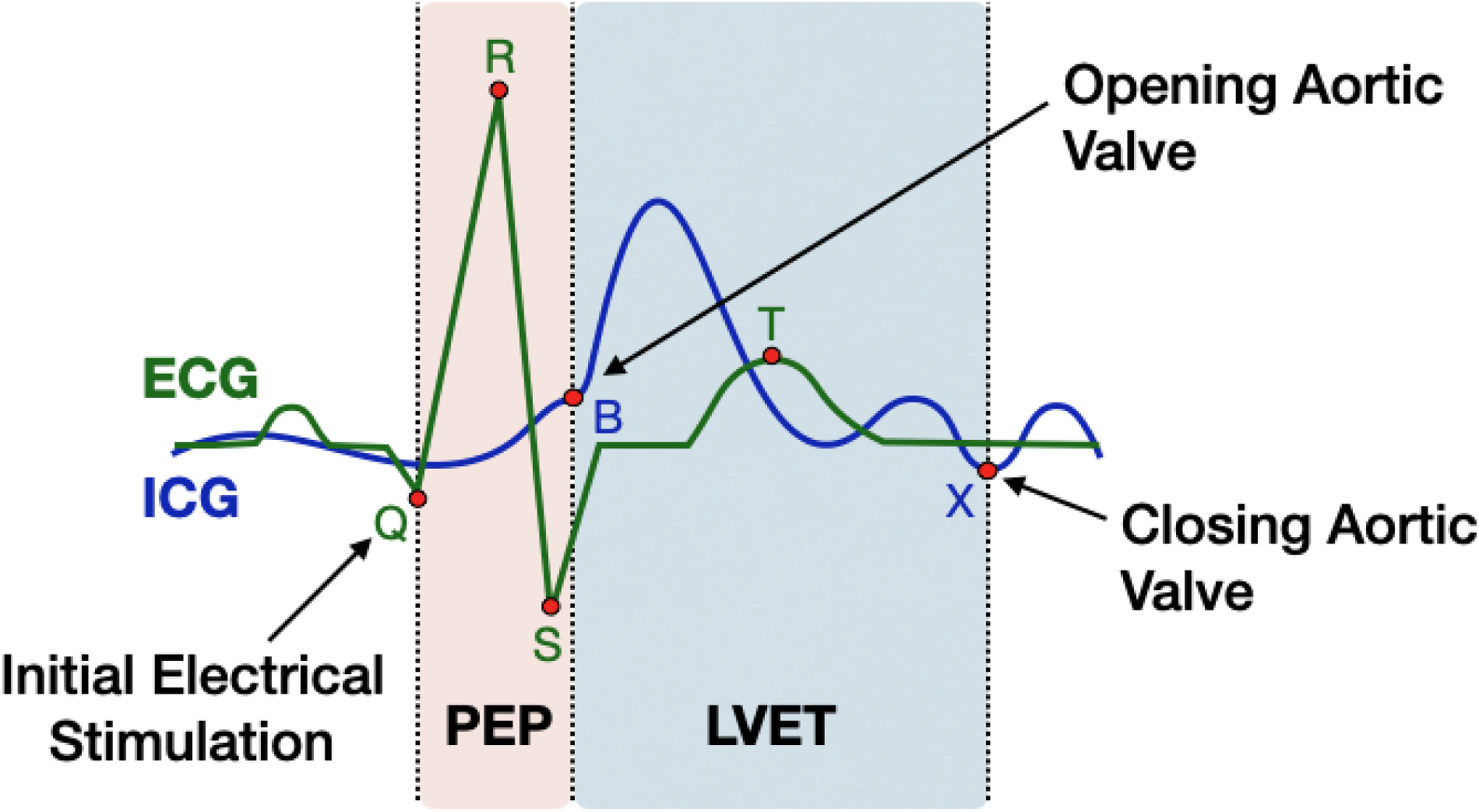
PEP and LVET measures extracted from ICG and ECG waveforms. The characteristic ICG waveform (in blue) displayed overlapping the ECG waveform (in green) with the ICG B-point representative of the opening of the aortic valve and the X-point as the closing of the aortic valve. The time interval between the Q-point of the ECG and the B-point of the ICG represents PEP the sum of electromechanical delay and isovolumic contraction of the ventricle. The time interval between b and x of ICG represents LVET.

Current methods for recording ECG and ICG are generally time intensive for researchers. In terms of apparatus preparation, the combined recording requires a total of 10 electrodes, placed on the neck, chest, and abdomen of the participant (Figure 2a). This procedure routinely requires ∼20 minutes and there can be significant variability in electrode placement by different researchers. There is also the issue of data processing. Current analysis techniques of the resulting time series have improved significantly with automated pipelines (Cieslak et al., 2018). Nevertheless, even with these software tools it can be difficult to localize precisely the opening of the aortic valve (the B-point) within the ICG signal. These software tools require the experimenter to build a classifier based off of 20 or more hand labelled B-points drawn from the full time series of data. For combined ICG-MRI experiments, the location of B-points is even more challenging and time consuming; 100 or more points must be manually labelled for classifiers to model an extended time series of data (Cieslak et al., 2018). Clearly, any technique that requires hand-labeling of the B-point, even if for just a subset of heartbeats, runs the risk of experimenter bias.

**FIGURE 2.**
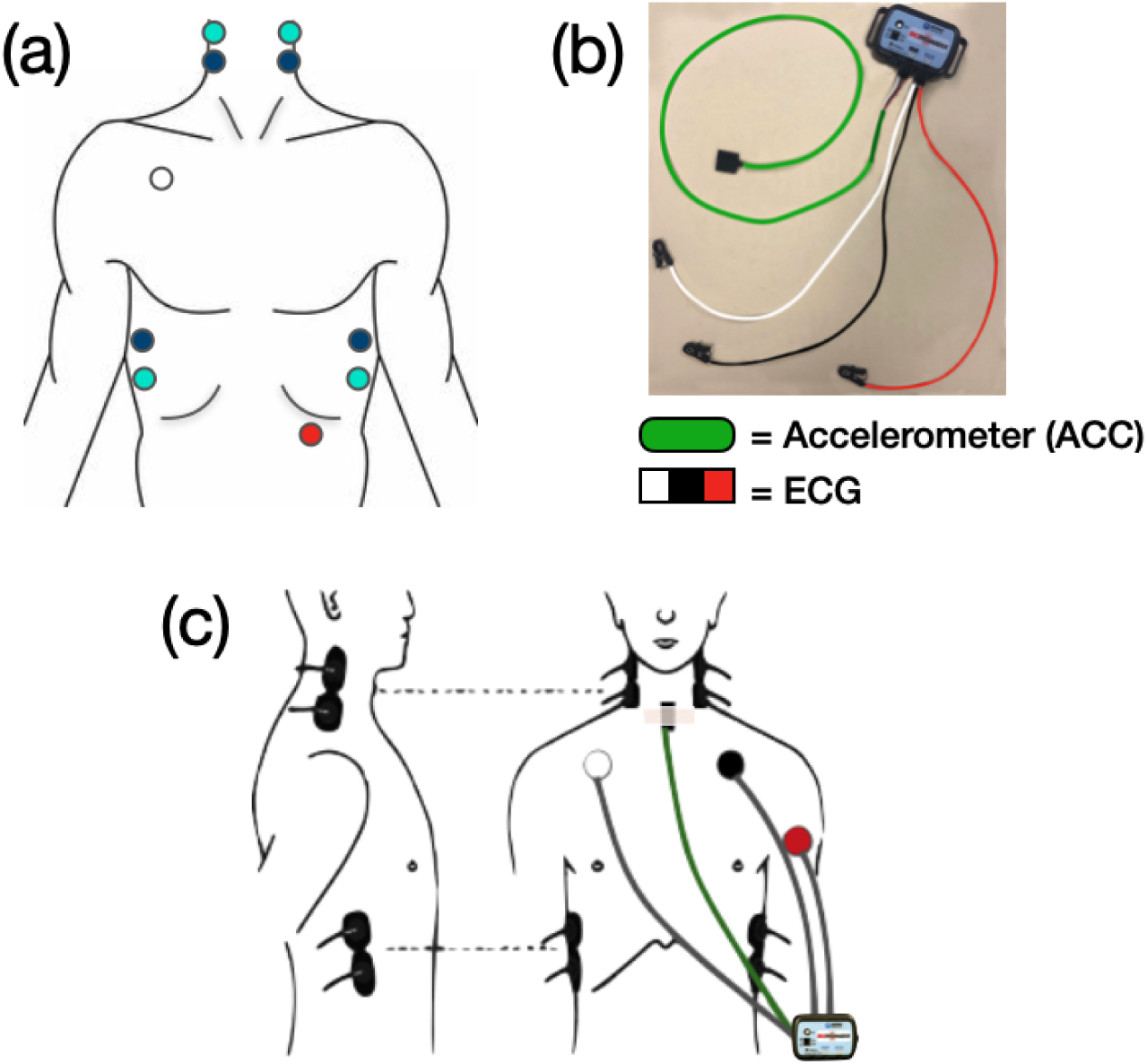
ICG and ECG setup and the Accelerometer Device (a) Original ICG and ECG setup uses a total of 10 electrodes. Eight ICG electrodes shown in blue: two on each side of the neck and two on each side of the torso. ICG electrodes in cyan are current-injecting electrodes, while electrodes in dark blue are voltage sensing. These are paired with two total ECG electrodes: one placed underneath the right collar bone (in white), and one placed underneath the left rib cage (shown in red). (b) A Biopac Inc. BioNomadix wireless transducer with three leads for ECG (represented by white, black, and red) and a single accelerometer (in green). (c) Experimental setup of the eight ICG electrodes (in black on either side of the neck and ribs), three ECG electrodes (white lead placed under right collarbone, black lead under the left collarbone, and red lead on the left tricep), and the ACC sensor (black rectangle secured in place with body tape at the suprasternal notch). ECG and ACC sensors are all connected to the BioNomadix wireless transducer.

To address these challenges in subject preparation and analysis, the authors worked with Biopac Systems Inc. to create a simple wearable and wireless heart monitor that includes 3-lead ECG, but replaces the 8 electrodes needed for ICG with a single-lead accelerometer (ACC) (Figure 2b). The ACC is placed flat in the suprasternal notch of the neck. Here, it senses motion orthogonal to the skin surface associated with the abrupt onset and offset of blood flow in the region of the aortic arch and great vessels. The resulting ACC waveform is composed of two smooth waves identifying key physiological events for quantifying both PEP and LVET, where the first peak represents the opening of the aortic valve, and the second peak represents its closing.

In this paper, we find that the ACC device can cut down on the time necessary for subject preparation and provide wave forms that simplify the subsequent data analysis while improving reliability. We also compare ACC based measures of PEP and LVET with measures from conventional ICG, and demonstrate equivalence between the two apparatuses, both in terms of event-related moment-to-moment fluctuations, and also in their estimation of the latency of key events resulting from a well-known and strong physiological perturbation – the Valsalva maneuver.

## 2 METHODS

### 2.1 Participants

Twenty healthy young adults (12 females, average age 23.5 years) were recruited to participate in this 1.5 hour study, for which they were compensated US $10/hour. All participants partook in a stressor condition where they performed the Valsalva maneuver. 19 participants performed a two-minute seated baseline session (one participant was unable to perform the seated baseline due to timing constraints). Following the first 8 participants, experimenters were given the opportunity to add a third session, in which the remaining 12 participants partook in a two-minute supine baseline session. All participants provided informed consent in accordance with the Institutional Review Board/Human Subjects Committee, University of California, Santa Barbara. They passed a screening protocol for physiological recording experiments to exclude anyone with a cardiovascular abnormality.

### 2.2 General procedure

Upon arrival, a research assistant described the general procedure of the study. Participants then completed consent forms and the screening form of cardiovascular related disease. Participants were taken into a private room where they changed into a surgical scrub top. A trained female research assistant placed 8 ICG electrodes on their neck and torso, and 3 ECG electrodes on the chest and upper left arm. Experimenters found that cleaning and exfoliating each area of the skin where electrodes would be placed was helpful for minimizing signal noise. Prior to placement of each electrode, an approximate 1-inch area of the skin was disinfected and exfoliated gently with an abrasive pad, followed by the application of Nu-Prep gel (ELPREP, Biopac Inc.), a skin exfoliant. Once the area was fanned dry, a small dab of electrode gel (GEL100, Biopac, Inc.) was placed on each of the 11 strong adhesive disposable foam electrodes (EL500, Biopac, Inc.) before they were placed on the body. For ICG (Figure 2c, in black), a total of 8 electrodes were placed on the torso and neck: two on each side of the neck, and two on each side of the torso as suggested by Bernstein (1986). Electrodes on the upper neck and lower torso (Figure 2a, in cyan) were each injecting a 4 mA alternating current into the thoracic cavity at 50 kHz, while the inner electrodes (Figure 2a, in blue) were voltage sensing. In combination, these electrodes provide basal trans-thoracic impedance (Z0) data and the first derivative (dZ/dt) of the pulsatile changes in transthoracic impedance. Inter-electrode distances between each pair of inner electrodes (Figure 2a, in cyan) were measured and recorded for analytic purposes. A total of three electrodes were used for ECG (Figure 2c, in white, black and red), one placed just under the right collarbone, one under the left collar bone, and one on the upper left arm. ICG electrodes provided the necessary grounding.

Participants were then taken to the experiment room and the ICG leads were connected to the respective hardware by carbon fiber leads. The ACC sensor was wiped down with an alcohol pad and allowed to dry between each participant. It was placed as flat on the suprasternal notch of the neck as possible with the wire oriented vertically and secured with 2-3 pieces of body tape (Figure 2c). Placement of all leads/sensors was standardized across participants and tasks. The ACC and ECG leads were permanently attached to an integrated wireless transducer that was placed in the pocket of the scrub shirt. During cardiovascular setup, the research assistant reviewed the importance of minimizing any and all movements of the seated posture throughout the course of the study in order to reduce noise in the cardiovascular readings. Research assistants estimated the time required to prepare each subject for the ACC method, from the start of skin exfoliation until the placement of the third ECG electrode. They added an estimate of the remainder of the time taken to apply the ICG electrodes, up until the recording of the inter-electrode ICG measurement, with the ACC time to produce an overall ICG preparation estimate. A minute was added to the ACC estimate to account for it being secured to the neck.

### 2.3 Cardiovascular equipment

Non-invasive physiological recording equipment was used for continuous real-time data acquisition throughout the study. All physiology equipment and software was from BIOPAC Systems, Inc. (Goleta, CA). Signals were integrated using the MP150 and collected at 2 kHz sampling rate. The ACC and ECG data were collected using a modified BN-RSPEC BioNomadix Transmitter to create a new “Cardio-Seismic” wireless transmitter unit. This unit employed a detachable +/- 2G accelerometer containing very low noise (20uG/sqrt(Hz)). The respiration input channel was modified to accommodate the accelerometer, while the ECG channel was left as is. ACC raw data was filtered from 0.1Hz to 100Hz. The ECG channel was filtered in real time from 1Hz-35Hz (the default bandwidth setting for that amplifier). However, additional filters were added using AcqKnowledge later in the processing chain (detailed under “ACC Signal Filtering”). ICG was collected using a NICO100C amplifier, ECG was collected using an ECG100C amplifier, and ACC was paired with an BN-RSPEC receiver. The ECG100C amplifier settings were adjusted to filter the incoming signal at 0.05Hz-35Hz and NICO100C amplifier settings were adjusted to filter at DC (no filter) - 10Hz. The BN-SPEC-R amplifier (ECG channel) was filtered in real-time from 1HZ-35Hz. These settings were kept constant across the sessions and therefore any delay contributions would be stable. All signals were displayed and stored on a laptop running AcqKnowledge software version 5.0.2.

### 2.4 Experimental protocol

To test the ACC’s reliability, a validation study was completed with a 15 second Valsalva maneuver, a classic physiological stressor known to trigger a complex cascade of autonomic reflexes, with an initial parasympathetic bradycardia and increase in peripheral sympathetic tone, followed by increased cardiac contractility that is based on sympathetic related drive (Blackburn et al., 1973; Ermishkin et al., 2007; Novak, 2011). Each experimental condition began with a two-minute baseline recording of ICG, ECG and ACC time series as the participants were in a seated posture with their hands resting on their thighs, breathing regularly. For each Valsalva trial, the participant was seated in a chair in a standard resting posture with their hands resting on their thighs. After a one minute baseline, a research assistant instructed them to begin the Valsalva maneuver. The instructions for this maneuver were standardized between participant to “bear down” on their abdominal muscles as if they were lifting something very heavy, creating a large amount of pressure in their gut region, and to hold this maneuver until the experimenter told them to release. Prior to this, each participant was instructed to do their best to remain in the same posture and to refrain as much as possible from any additional movement. After 15 seconds, the research assistant informed the participant that they may release the Valsalva and return to regular breathing for a two-minute recovery phase to allow autonomic activity to return to baseline. We had the opportunity to run 12 of the participants in an additional two-minute condition in a supine position, lying flat on a gurney with their arms at their sides in the same experiment room.

### 2.5 ACC signal filtering

An initial finite impulse response (FIR) bandpass filter of 20 to 30 Hz was applied to the ACC signal. The signal was then converted to its absolute value, after which an additional low pass filter of 15 Hz was applied. Finally, mean-value smoothing was applied at a smoothing factor of 221 samples. All these filters were applied post-collection and pre-processing. Following these filters, experimenters were able to convert the original ACC signal into two smooth waves, with the first wave’s peak indicating the opening of the aortic valve, and the second wave’s peak indicating the closing (Figure 3a). Collectively, these measures are capable of potentially replacing the entire ICG setup, as the main points necessary to measure the SNS and other ANS perturbations are the mechanistic movement of the aortic valves (Figure 3b).

**FIGURE 3.**
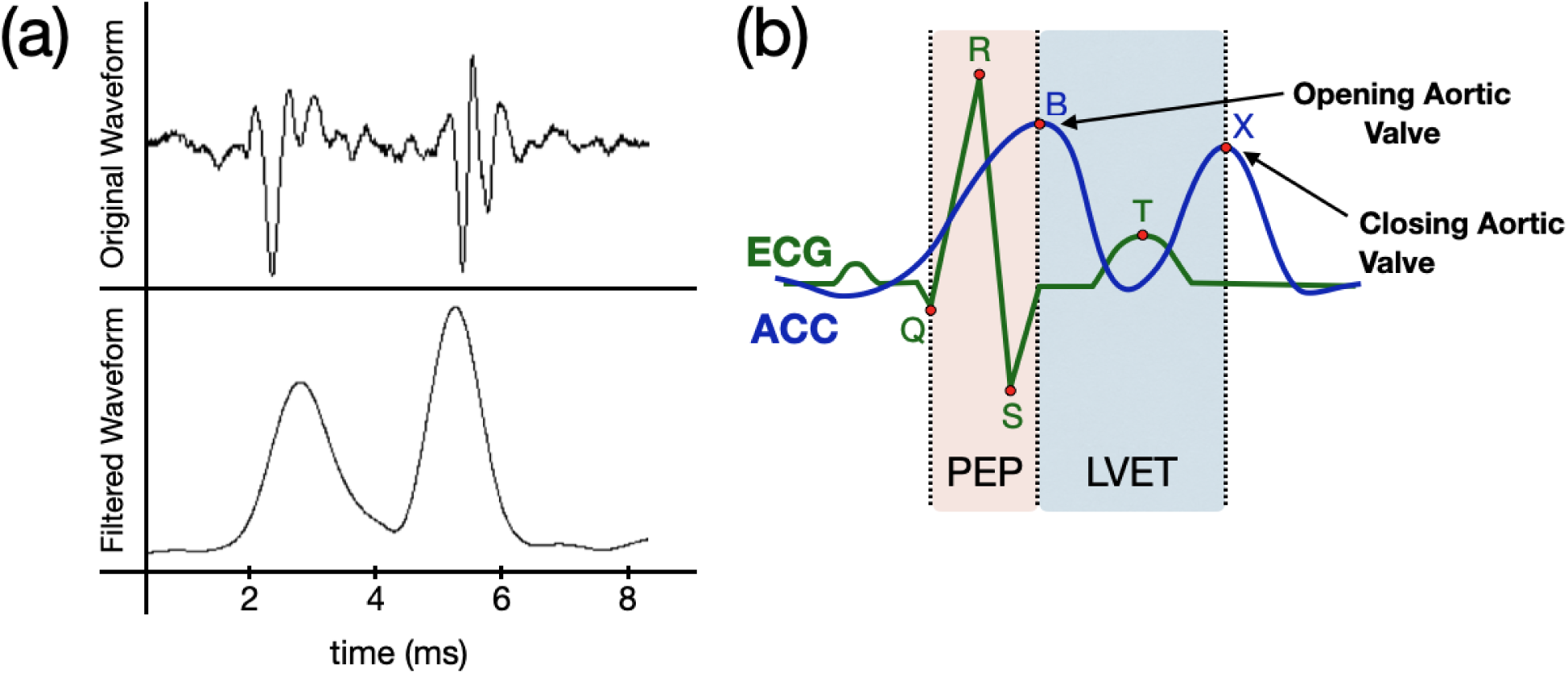
Extracting PEP and LVET using characteristic ECG and filtered ACC waveforms. (b) Original ACC waveform of a single heartbeat representing movement in the z-direction of the mechanistic pulse wave moving through the trachea, displayed above the filtered waveform. The first peak of the filtered waveform is analyzed as the opening of the aortic valve (the B-point). The 2^nd^ peak is analyzed as the closing (the X-point). (b) The filtered ACC waveform (in blue) is displayed overlapping the ECG waveform (in green). The peak of the first waveform of the ACC is labeled as the B-point (opening of the aortic valve), while the peak of the second waveform is labeled as the X-point (closing of the aortic valve). The time interval between the Q-point of the ECG waveform and B-point peak of the ACC is measured as PEP. The time interval between the b- and x-peak of the ACC is measured as LVET.

### 2.6 Cardiovascular analysis

Analysis of the ICG time series, including preprocessing and B-point and X-point labeling using the open-source Moving Ensemble Analysis Pipeline (MEAP) has been described in detail previously (Cieslak et al., 2018). The MEAP software was modified to also import the filtered ACC time series where automated B- and X-point labels were applied to the two peaks of the ACC signal. Trained research assistants manually checked each classification and edited out any artifacts in the ICG and ACC time series. The automation of ACC time point classifications required little to no outlier edits, while the same time point estimations for ICG required significant outlier detection and hand labeling. For this study, the ICG and ACC data were each combined with the ECG times series and processed with a moving ensemble-average along a 15 second window, allowing for continuous estimations of two physiological measures of interest: PEP and LVET. Research assistants estimated the time required to perform manual checking of standard ICG and ACC time series. For ACC, this time was estimated from the start of opening the file in MEAP, up until the completion of the correction of any obvious outliers following the initial computation of moving ensembles. Following this step, the research assistant created an ICG B-point classifier based off 25 hand-labeled B-points, applied this classifier to the data, and repeated the correction for outliers. This remaining time was added to the previous ACC estimate to create an ICG analysis estimate. To control for reliability across research assistants scoring the data, the same research assistant analyzed within participant trials.

### 2.7 Statistical analyses

To compare the ICG and ACC derived physiology time series we conducted four analyses. The first analysis estimated the overall similarities between moment to moment fluctuations recorded by both sensors. We computed a summarized delta time series for each measure: change in average activity across successive five second windows. For both PEP and LVET, and in each of the seated, supine and Valsalva conditions, we tested the similarities between the ICG and ACC delta time series across subjects. To this end, we used a linear mixed effects (random intercept) model, that tested for agreement between moment-to-moment changes in our physiological indices (PEP, LVET) recorded with ICG and ACC. Given the relatively small size of this exploratory sample, we also included a hierarchical Bayesian analogue of the linear mixed effects model (hierarchical regression), to directly capture the uncertainty of our estimated model parameters in a manner that accounts for between subject variability.

In the second analysis we assessed whether both sensors would estimate similar latency of event-related peak physiological change during our Valsalva stressor condition. For this, we used a hierarchical Bayesian changepoint model; the model predicts the likelihood that the summarized delta time series have two means. From these two distributions a switchpoint can be identified. In theory, one side of the peak should be predominantly unidirectional and a mean of one sign, while the return to baseline should be unidirectional in the opposite direction. The switchpoint therefore estimates the critical point when there is a maximal change between distributions, without assuming signal valence. If the ACC is similar to ICG then this switchpoint should be at the same time.

In the third and fourth analyses, we examined the internal consistency (i.e., within session reliability) and criterion validity (i.e., cross-instrument), separately for each subject, and for PEP and LVET values expressing the percentage change in the Valsalva condition, relative to the baseline condition at each data point. To align baseline and Valsalva conditions for meaningful percentage-change estimation, all sessions were cropped to the first 120 seconds worth of data and linearly resampled to provide one estimate of physiology (PEP or LVET) at each second.

All linear mixed effects models were fitted using the lmerTest (Kuznetsova et al., 2017) package in R (R Core Team, 2020). Bayesian hierarchical regression, changepoint, internal consistency and criterion validity analyses were run using PyMC3 (Salvatier et al., 2016) libraries in Python 3. All other data preprocessing was carried out in MATLAB version 2020a.

## 3 RESULTS

We found that the ACC method cut down on human labor estimates from about 20 minutes for application/preparation and 1 hour for analysis per 1 hour of data collected, to about 5 minutes of application and 20 minutes of analysis per hour of data. To compare the results between the performance of the ACC and ICG, we examined the time courses of two temporal features (PEP and LVET), using our two different methods (the ACC and ICG). The following analyses were used to determine the validity of the ACC compared to the classic ICG method for both PEP and LVET measures. Mean and standard deviation for both ACC and ICG-derived PEP and LVET are summarized in tables 1 and 2. (To show individual participant consistency between the two measures, Bland-Altman plots between the ICG and ACC delta timeseries of PEP and LVET scores was performed. To assess whether the ACC can also model respirations, a power coherence analysis comparing respiration between ACC and ICG was calculated. Additionally, to examine whether our estimates of PEP and LVET relate similarly to another physiological measure, we performed comparisons of ACC and ICG derived heart rate variability with PEP and LVET. The results of these three analyses are available in Supplementary Materials.)

1. *Linear mixed effects (random intercept) model.* We tested the similarities between the ICG and ACC delta time series across participants. For instance, as ICG’s PEP values change, does the ACC’s PEP change with it? We fitted separate models for PEP and LVET, in each case modeling the signals measured by ICG as a function of the signals measured from ACC. The model fitted a single parameter ACC and a separate intercept for each subject (random intercept model).
2. *Hierarchical Bayesian delta regression.* The concern with the random intercepts model (which only fits a single ACC beta to account for all subjects) is that significant effects may be driven by a single outlier participant. To mitigate the effects of individual participants, we ran a Bayesian regression as a group-based analysis, which estimates the parameters (mu and sigma) of a hierarchical Gaussian distribution from which each subject’s ACC beta is drawn. Significance at the group level is determined by the posterior distribution of the hierarchical distribution’s mu parameter exceeding 0, using a 94% highest density interval (HDI). In other words, allowing for individual differences, the mean of the distribution that characterizes betas across all subjects is confidently above or below 0.
3. *Hierarchical Bayesian changepoint in delta time course.* While the previous two analyses were performed for all three trials (seated baseline, supine, and Valsalva), this changepoint analysis was only performed on the Valsalva condition. For these Valsalva trials, we assume that the physiologic measure (either PEP or LVET) will undergo a significant change at some time point after the initiation of the maneuver. Thus, we assume deltas in the time series of each measure can be best categorized by two distributions, one distribution for a change in one direction (e.g., with a positive mean delta if measure increases) and another, with an opposite signed mean when the measure returns to baseline. We also estimate the changepoint, i.e., the point in time that most likely reflects when the observed deltas transition from being best summarized by one distribution to the other. In other words, this model determines when the critical point of an event-related change in deltas occurs in both LVET and PEP (peak max or peak min) in response to the Valsalva maneuver, without a priori assumptions on the direction of change. This analysis was performed only on the Valsalva trials.
4. *Internal Consistency.* We measured the consistency of each subject’s data recorded across discrete moments in time. For each of the eight measurements - two conditions (BL,V), two sensors (ACC, ICG) and two physiology measures (PEP, LVET) - we computed a second-by-second estimate of the standard deviation (SD) of the distribution for that measurement across subjects. We fitted these SD values using a Bayesian model. By computing a posterior for each SD, we could compare the SD at every timepoint, with the SD for each other time point for a given measurement. The results confirm very strong internal consistency for all measurements. Within each of the eight measurements, we saw no strong evidence that the SD across subjects at any timepoint was credibly different to those at all other timepoints (See Supplementary Materials for an exhaustive pairwise departure analysis). We conclude that both ICG and ACC record consistent measures of both PEP and LVET across subjects at discrete moments in time, in both the baseline and Valsalva condition (Figures 10b and 11b).
5. *Criterion Validity.* We assessed criterion validity between departures from the baseline condition driven by the Valsalva condition. For both PEP and LVET, we computed the percentage difference between each timepoint of the Valsalva condition and the corresponding timepoint in the baseline condition (% Δ V-BL), for both ACC and ICG for each subject. We then used a Bayesian model to estimate the mean of the distribution of subjects’ (% Δ V-BL) at each second in time for both ICG and ACC. Sampling each timepoint’s posterior for mean (% Δ V-BL – ICG) and mean (% Δ V-BL – ACC), we computed a distribution of Pearson coefficients that quantified the relation between ICG Valsalva-baseline departure and ACC Valsalva-baseline departure across 120 seconds of data. For both PEP and LVET, this n=10,000 simulation produced a distribution of all positive coefficients, confirming a strong association between measurements derived by the ACC and ICG, using a procedure that accounted for the random effects at each timepoint due to individual difference (Figures 10a and 11a).

**TABLE 1.**
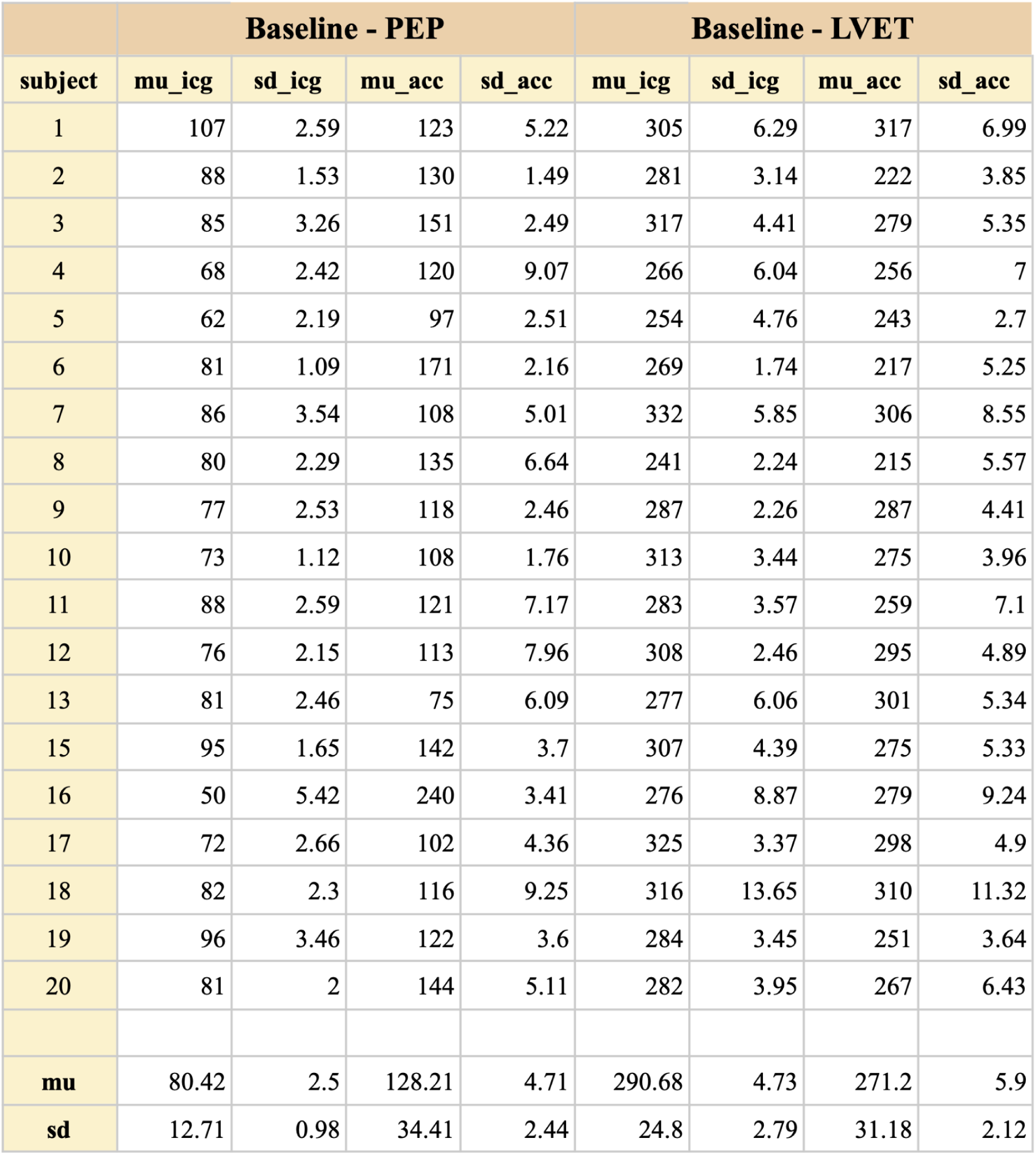
Summary data for PEP and LVET for the baseline condition. Mean (mu) and standard deviations (sd) for both ACC and ICG-derived PEP and LVET (across the entire session). Participant 14 did not have a BL condition.

**TABLE 2.**
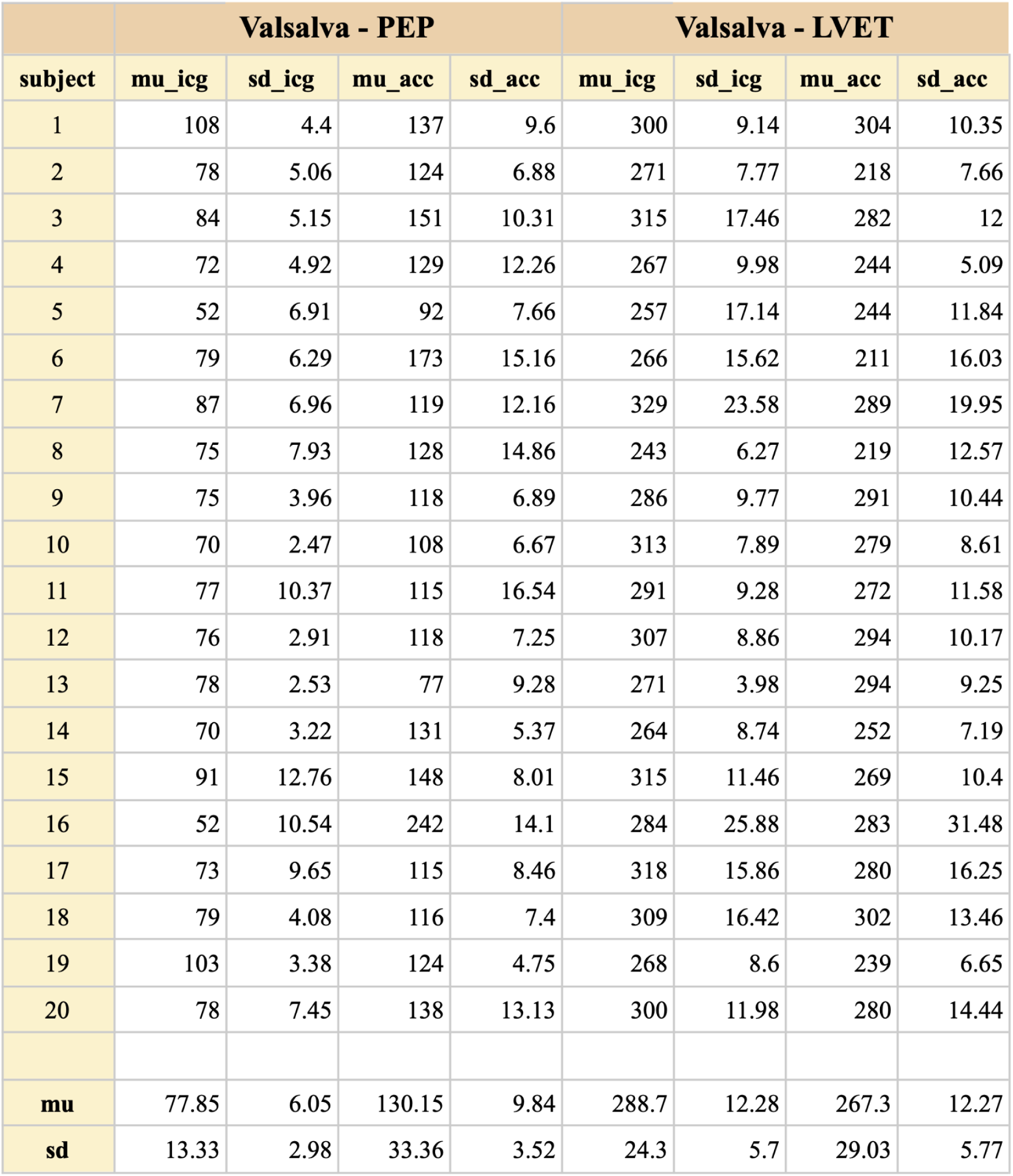
Summary data for PEP and LVET for the Valsalva condition. Mean (mu) and standard deviations (sd) for both ACC and ICG-derived PEP and LVET (across the entire session).

### 3.1 Pre-ejection period (PEP)

Overall, PEP measures collected using the ACC closely followed PEP measures from ICG. Results of the linear mixed effects model for 19 participants in a seated two-minute baseline confirmed a positive relationship between the delta of the ACC’s PEP with the ICG’s PEP (β = 0.216, SE = 0.059, p < 0.001). In the Bayesian delta regression of the same baseline period, the mean of the hierarchical posterior distribution of subjects’ beta values was 0.102 with a 94% HDI of -0.012, 0.218], indicating a marginally positive relationship between ACC and ICG recorded PEP perturbations (Figure 4).

**FIGURE 4.**
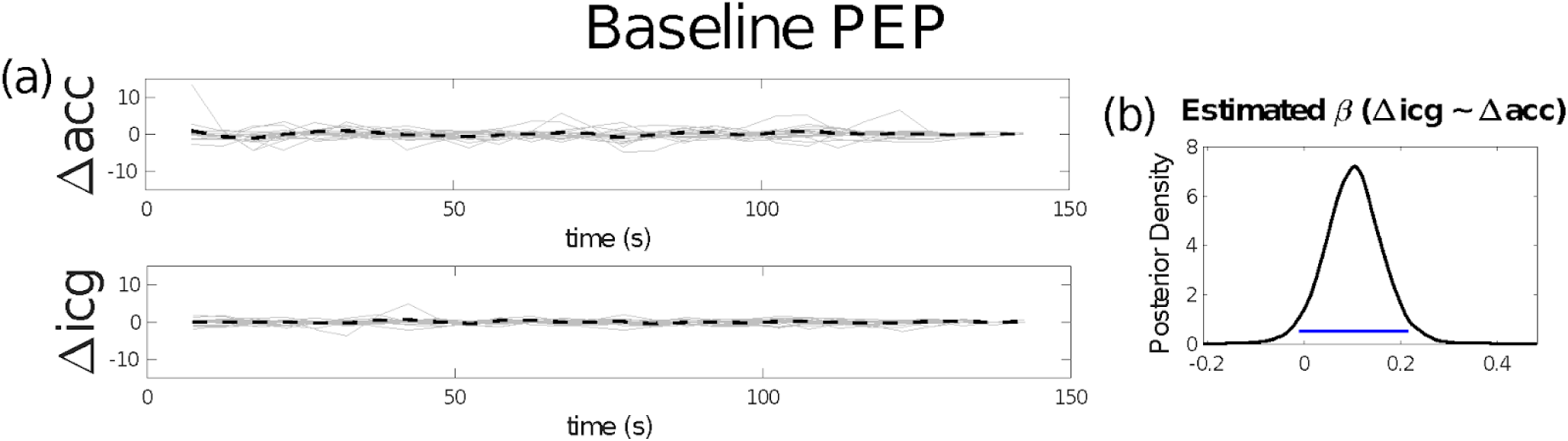
Pre-ejection period two-minute seated baseline hierarchical Bayesian delta regression model. (a) Summarized delta time series for each of ACC’s and ICG’s PEP, each time point reflects the change (delta) in average activity across successive five second windows. Light gray lines represent data from individuals, while dashed black lines represent the central tendency. (b) A Bayesian estimate of the mean of a Gaussian distribution from which each subject’s beta parameter (modeling ICG as a function of ACC) is drawn. The blue line denotes the 94% Highest Density Interval (HDI). Most of the HDI above 0 reflects a marginally positive relationship between the deltas estimated with ICG and ACC.

Results of the linear mixed effects model for 12 participants in a supine two-minute baseline confirmed a positive relationship between the delta of the ACC’s PEP with the ICG’s PEP (β = 0.216, SE = 0.059, p < 0.001). In the Bayesian delta regression of the same baseline period, the mean of the hierarchical posterior distribution of subjects’ beta values was 0.079 with a 94% HDI of [-0.016, 0.182], indicating a marginally positive relationship between the ACC and ICG recorded PEP perturbations (Figure 5).

**FIGURE 5.**
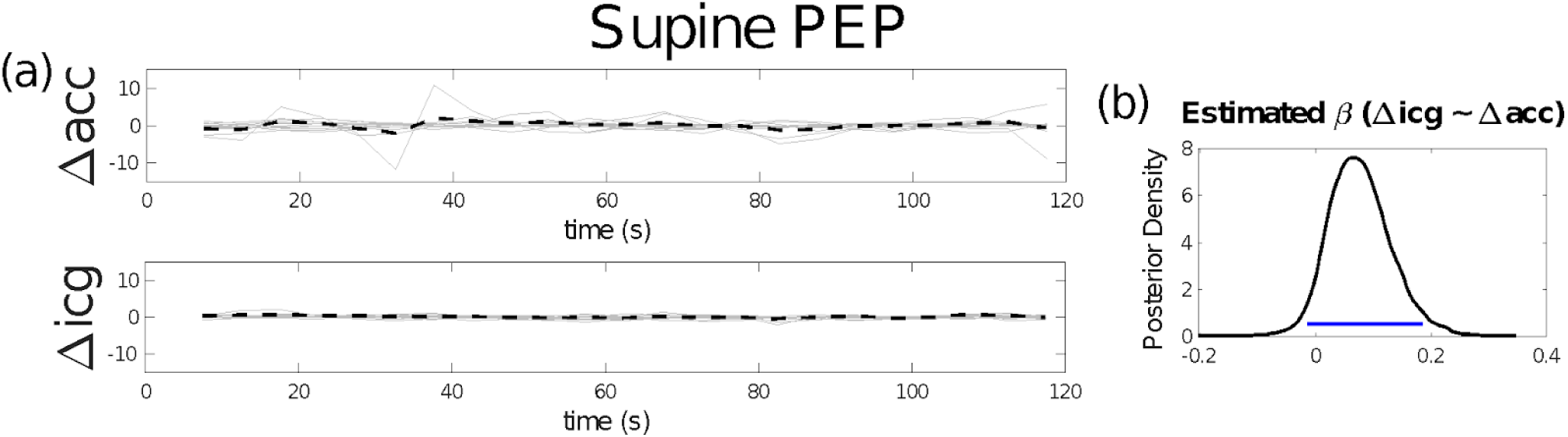
Pre-ejection period two-minute supine hierarchical Bayesian delta regression model. (a) Summarized delta time series for each of ACC’s and ICG’s PEP, each time point reflects the change (delta) in average activity across successive five second windows. Light gray lines represent data from individuals, while dashed black lines represent the central tendency. (b) A Bayesian estimate of the mean of a Gaussian distribution from which each subject’s beta parameter (modeling ICG as a function of ACC) is drawn. The blue line denotes the 94% Highest Density Interval (HDI). Most of the HDI above 0 reflects a positive relationship between the deltas estimated with ICG and ACC.

Results of the linear mixed effects model for 20 subjects performing Valsalva confirmed a positive relationship between the delta of the accelerometer’s PEP with the ICG’s PEP (β = 0.688, SE = 0.097 p < 0.001). Additionally, in the Bayesian delta regression during the Valsalva, the mean of the hierarchical posterior distribution was 0.168 with a 94% HDI of [0.076, 0.259], indicating a positive relationship between ACC and ICG recorded perturbations (Figure 6a). The Bayesian changepoint model estimated that peak ICG-recorded PEP change occurred at 82.79 seconds (posterior mean; note that the Valsalva onset was at 60 seconds), while the peak of ACC-recorded PEP change occurred at 80.75 seconds (posterior mean). In both cases, the changepoints characterizing these peaks marked the point in time where the delta of PEP went from a negative to a positive slope (Figure 6c), consistent with the expected increase in sympathetic drive following the Valsalva. Of note, the ICG’s changepoint 94% HDI [73.4, 92.3] fell 100% within the ACC’s changepoint 94% HDI [69.15, 92.35]. We therefore observe overall strong agreement between PEP measures recorded from the ACC and ICG in all trials (baseline, supine, and Valsalva), with the strongest positive relationship occurring in the Valsalva condition.

**FIGURE 6.**
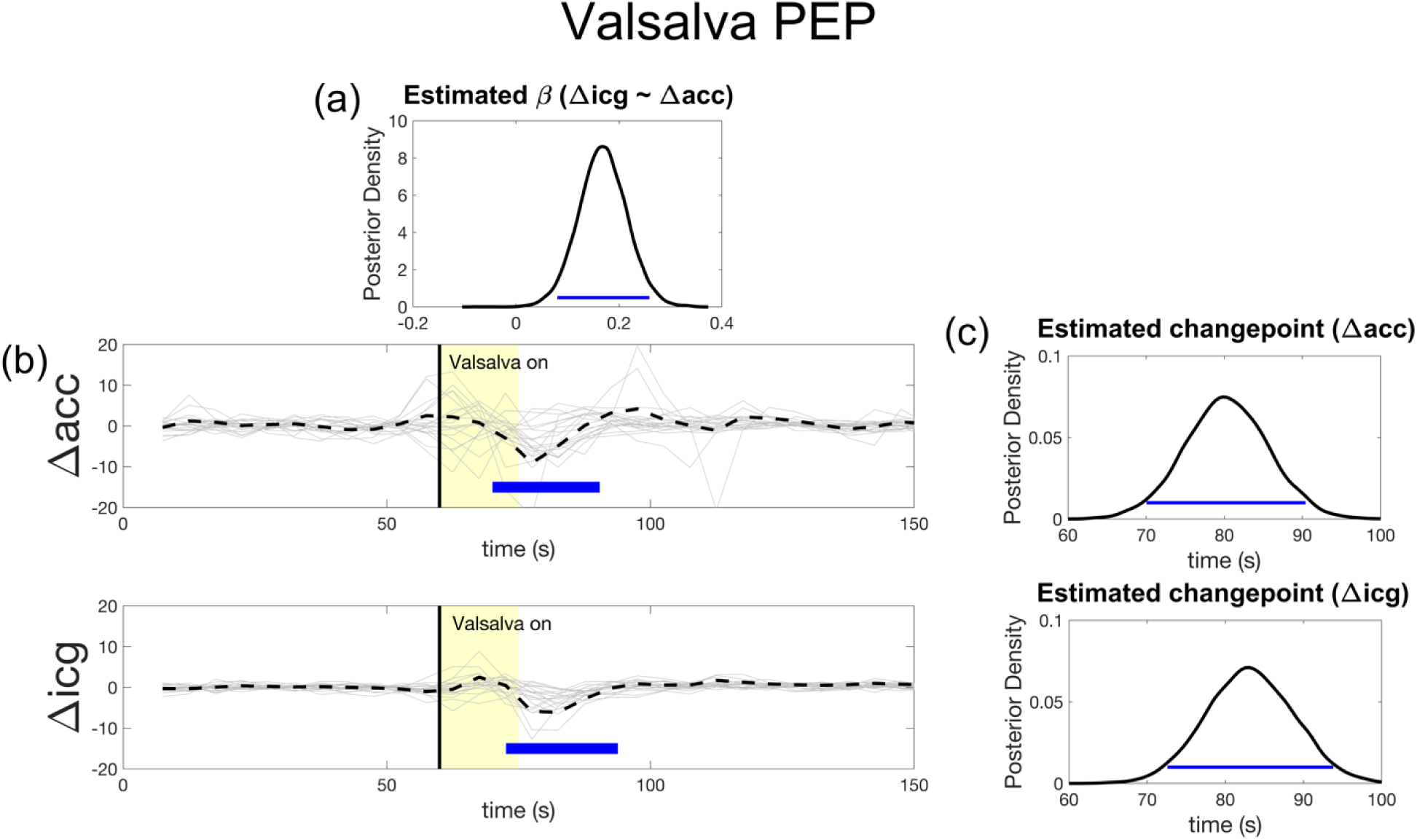
Valsalva pre-ejection period hierarchical Bayesian models. The blue line denotes the 94% Highest Density Interval (HDI). Most of the HDI above 0 reflects a positive result. (a) A Bayesian estimate of the mean of a Gaussian distribution from which each subject’s beta parameter (modeling ICG as a function of ACC) is drawn. (b) Summarized delta time series for each of ACC’s and ICG’s PEP, each time point reflects the change (delta) in average activity across successive five second windows. Light gray lines represent data from individuals, while dashed black lines represent the central tendency. (c) A Bayesian model that estimated the latency of peak event-related change driven by the Valsalva, separately for the ACC and ICG recorded PEP. The latency of this peak is the modeled changepoint between two distinct distributions of delta values (in this case mostly negative distribution and a mostly positive distribution).

### 3.2 Left ventricular ejection time (LVET)

Results of the linear mixed effects model for 19 participants during baseline recording confirmed a positive relationship between the delta of the ACC’s LVET with the ICG’s LVET (β = 0.885, SE = 0.118, p < 0.001). In the Bayesian delta regression of the LVET recorded during baseline, the mean of the hierarchical posterior distribution of subjects’ beta values was 0.349 with a 94% HDI of [0.213, 0.499], supporting a positive relationship between ACC and ICG recorded LVET perturbations during a seated baseline (Figure 7).

**FIGURE 7.**
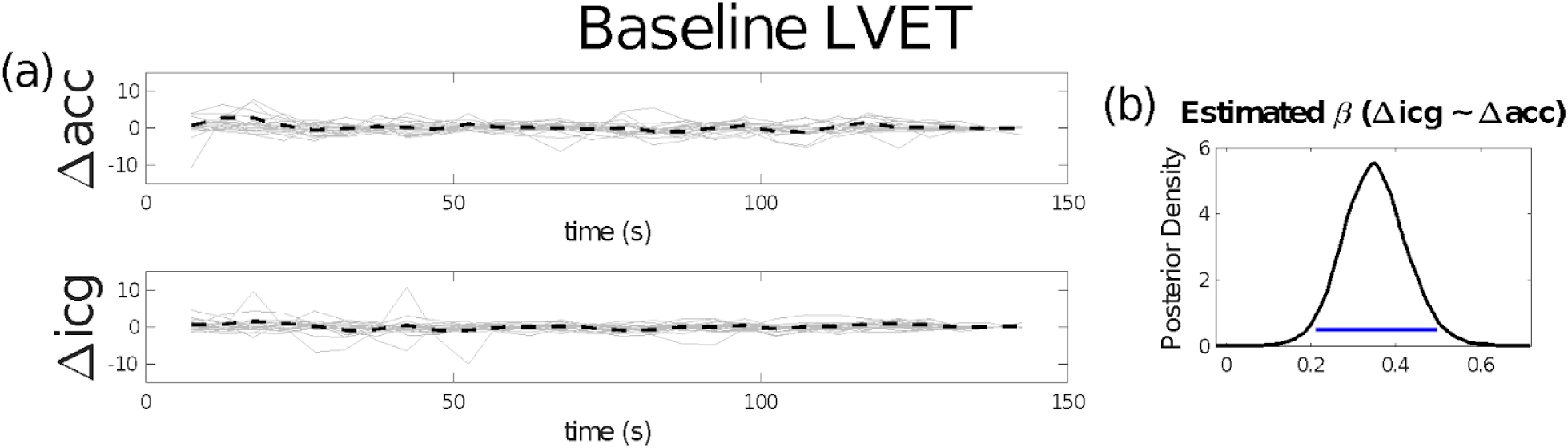
Left ventricular ejection time two-minute baseline hierarchical Bayesian delta regression model. a) Summarized delta time series for each of ACC’s and ICG’s LVET, each time point reflects the change (delta) in average activity across successive five second windows. Light gray lines represent data from individuals, while dashed black lines represent the central tendency. (b) A Bayesian estimate of the mean of a Gaussian distribution from which each subject’s beta parameter (modeling ICG as a function of ACC) is drawn. The blue line denotes the 94% Highest Density Interval (HDI). Most of the HDI above 0 reflects a positive relationship between the deltas estimated with ICG and ACC.

Results of the linear mixed effects model for 12 participants during supine recording showed no relationship between the delta of the ACC’s LVET with the ICG’s LVET (β = -0.014, SE = 0.1, p > 0.1). In the Bayesian delta regression of LVET recorded during the supine condition, the mean of the hierarchical posterior distribution of subjects’ beta values was 0.178 with a 94% HDI of [-0.021, 0.401], indicating a marginally positive relationship between the ACC and ICG recorded LVET perturbations while lying in a supine position (Figure 8).

**FIGURE 8.**
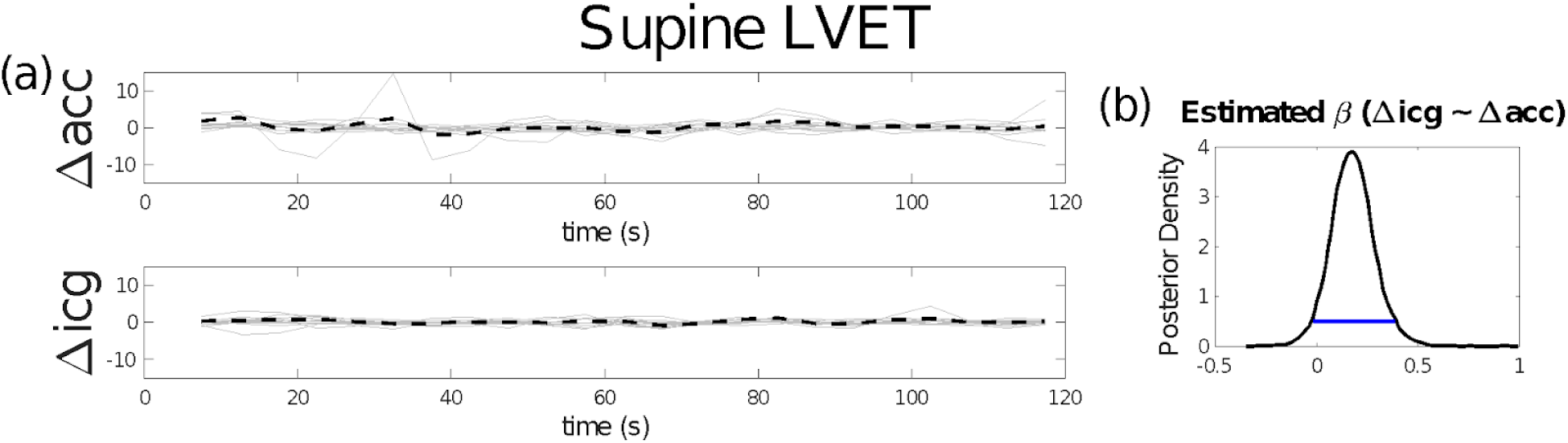
Left ventricular ejection time two-minute supine hierarchical Bayesian delta regression model. (a) Summarized delta time series for each of ACC’s and ICG’s LVET, each time point reflects the change (delta) in average activity across successive five second windows. Light gray lines represent data from individuals, while dashed black lines represent the central tendency. (b) A Bayesian estimate of the mean of a Gaussian distribution from which each subject’s beta parameter (modeling ICG as a function of ACC) is drawn. The blue line denotes the 94% Highest Density Interval (HDI). Most of the HDI above 0 reflects a positive relationship between the deltas estimated with ICG and ACC.

Results of the linear mixed effects model for 20 subjects performing Valsalva confirmed a positive relationship between the delta of the ACC’s LVET with the ICG’s LVET (β = 3.581, SE = 0.208, p < 0.001). In the Bayesian delta regression of LVET during Valsalva, the mean of the hierarchical posterior distribution of subjects’ beta values was 0.459 with a 94% HDI of [0.289, 0.631], supporting this positive relationship between ACC and ICG recorded LVET perturbations during a Valsalva (Figure 9a). The Bayesian changepoint model estimated that peak ICG-recorded LVET change occurred at 79.1 seconds (posterior mean; note that the Valsalva onset was at 60 seconds), while the peak of ACC-recorded LVET change occurred at 78.6 seconds (posterior mean). In both cases the changepoints characterizing these peaks marked the point in time where the delta of LVET went from a positive to a negative slope (Figure 9c), consistent with the expected increase in autonomic drive following the Valsalva. Of note, the 94% HDI of the ACC’s changepoint [69.1, 87.95] fell 94.15% within the HDI of the ICG’s changepoint [70.25, 91.55]. We therefore observe strong agreement between LVET measures recorded from the ACC and ICG in both baseline and Valsalva trials, with the strongest positive relationship occurring in the Valsalva condition.

**FIGURE 9.**
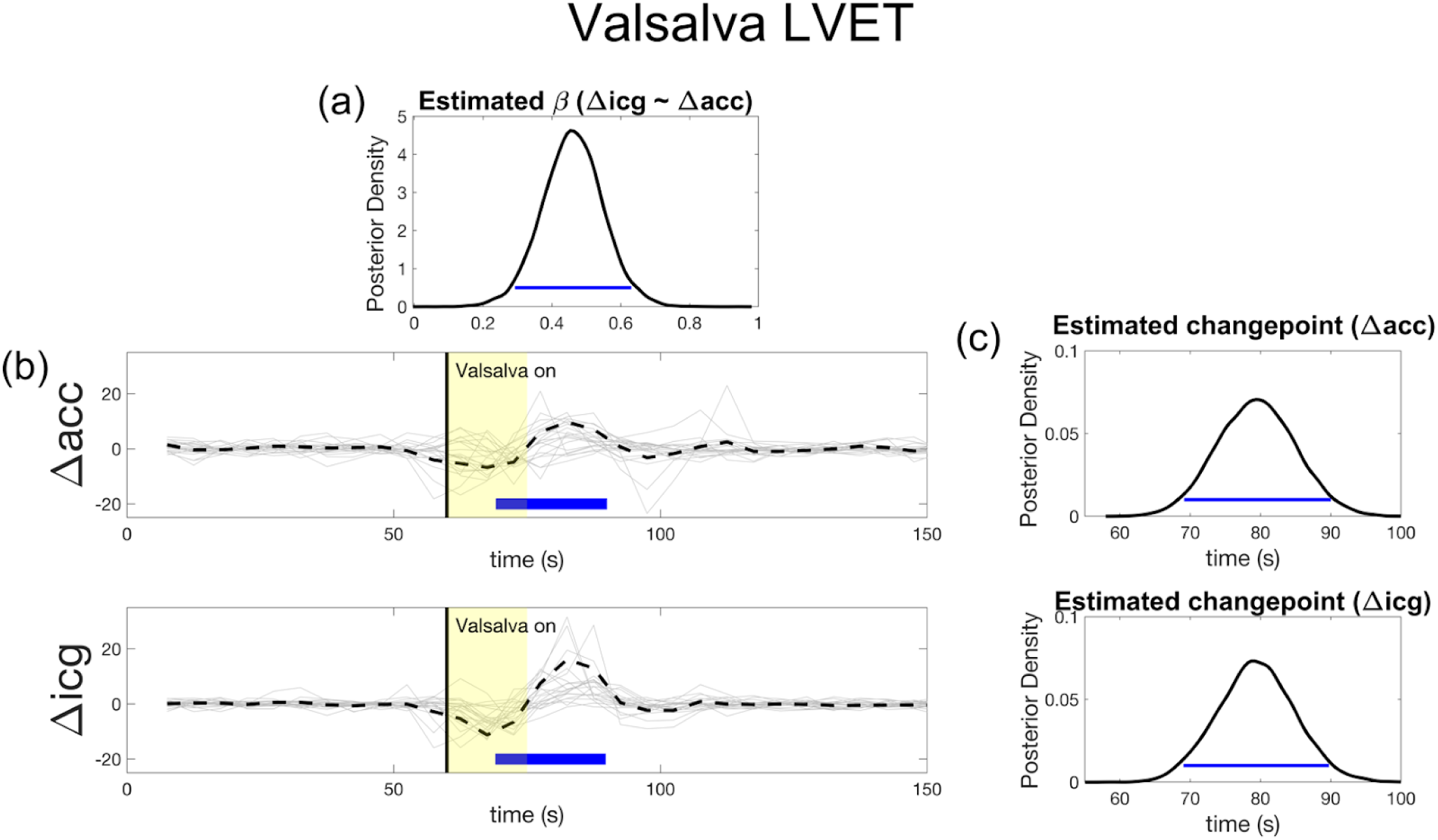
Valsalva left ventricular ejection time hierarchical Bayesian models. The blue line denotes the 94% Highest Density Interval (HDI). Most of the HDI above 0 reflects a positive result. (a) A Bayesian estimate of the mean of a Gaussian distribution from which each subject’s beta parameter (modeling ICG as a function of ACC) is drawn. (b) Summarized delta time series for each of ACC’s and ICG’s LVET, each time point reflects the change (delta) in average activity across successive five second windows. Light gray lines represent data from individuals, while dashed black lines represent the central tendency. (c) A Bayesian model that estimated the latency of peak event-related change driven by the Valsalva, separately for the ACC and ICG recorded LVET. The latency of this peak is the modeled changepoint between two distinct distributions of delta values (in this case mostly negative distribution and a mostly positive distribution).

**FIGURE 10.**
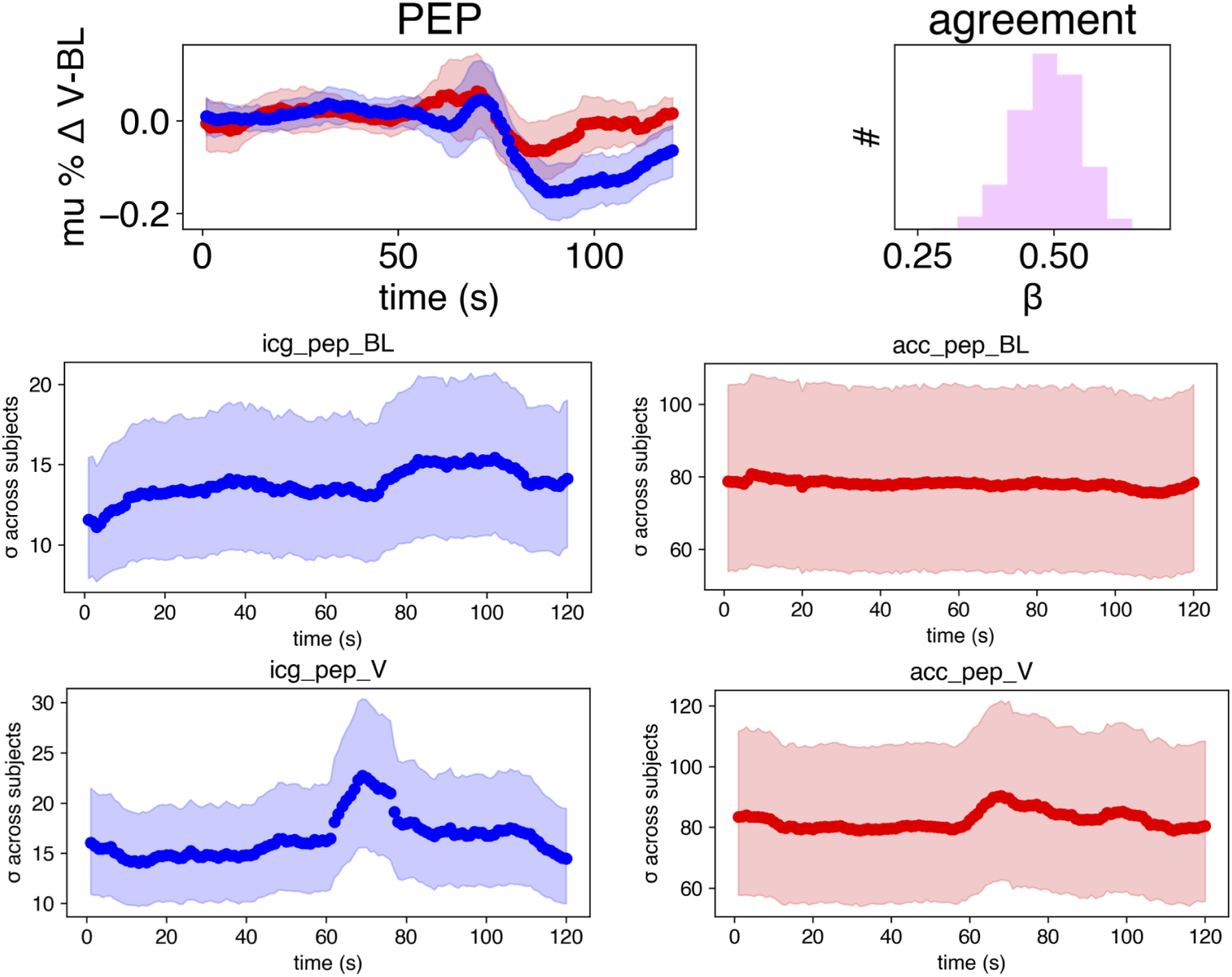
PEP internal consistency and criterion validity. (a) Criterion validity between departures from the baseline condition driven by the Valsalva condition (% Δ V-BL). For ICG (blue) and ACC (red) we estimated the mean (% Δ V-BL - dots) across subjects at each data point, with the shaded region reflecting the HDI of each mean estimate. Agreement histogram on the right represents the distribution of Pearson correlation coefficients from n=10,000 draws from the (% Δ V-BL - ACC) and (% Δ V-BL - ICG) posteriors at each data point. (b) Dots show the mean of the posterior estimating the standard deviation across subjects at each data point, while the shaded region depicts the HDI, i.e., the credible range of variability across subjects at each moment in time.

**FIGURE 11.**
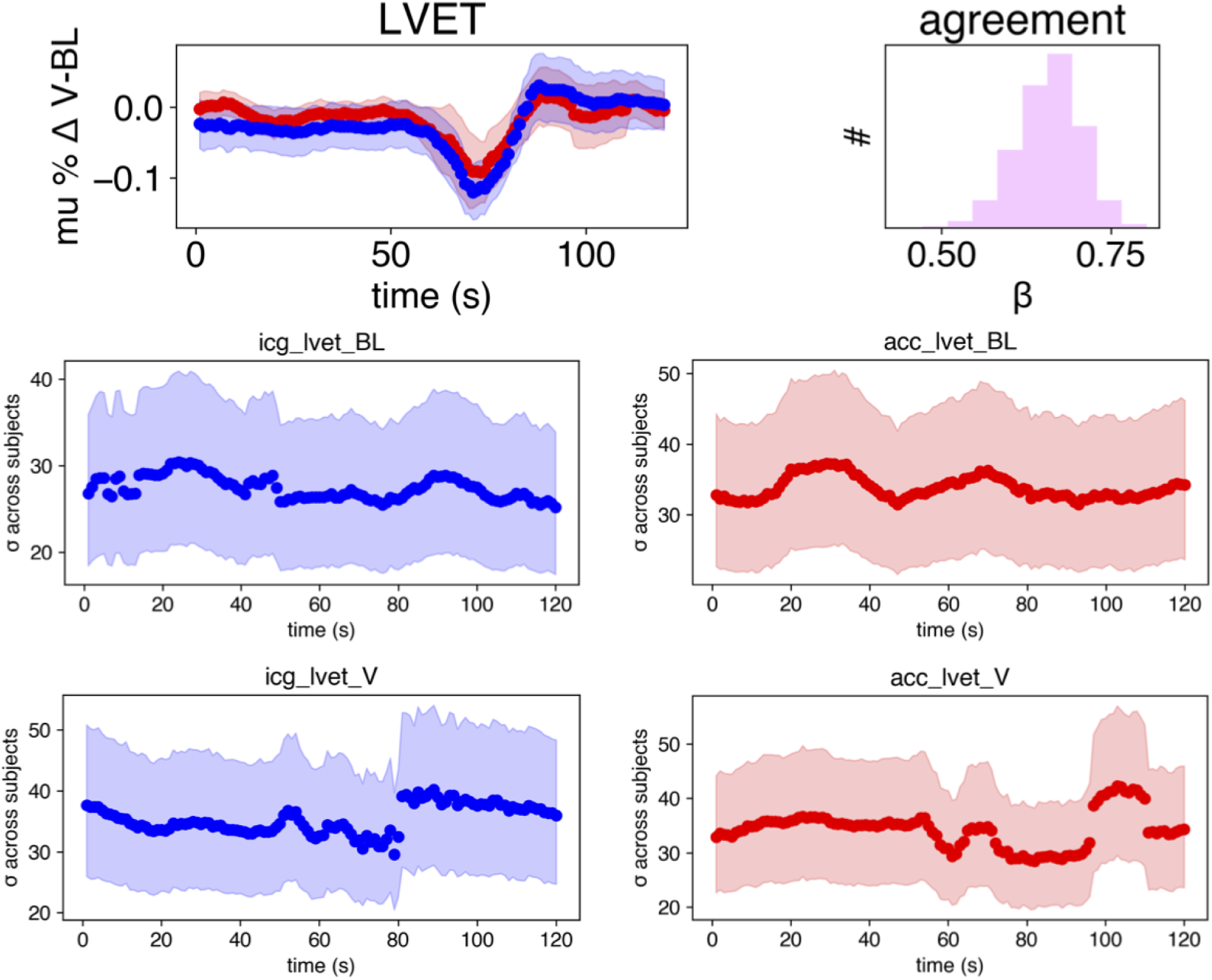
LVET internal consistency and criterion validity. (a) Criterion validity between departures from the baseline condition driven by the Valsalva condition (% Δ V-BL). For ICG (blue) and ACC (red) we estimated the mean (% Δ V-BL - dots) across subjects at each data point, with the shaded region reflecting the HDI of each mean estimate. Agreement histogram on the right represents the distribution of Pearson correlation coefficients from n=10,000 draws from the (% Δ V-BL - ACC) and (% Δ V-BL - ICG) posteriors at each data point. (b) Dots show the mean of the posterior estimating the standard deviation across subjects at each data point, while the shaded region depicts the HDI, i.e., the credible range of variability across subjects at each moment in time.

## 4 DISCUSSION

The use of ICG has been an incredibly useful and strong tool in measuring the autonomic nervous system, although it is not without its challenges. Current ICG methods suffer from time intensive preparation and analysis. The main goal of the present study was to investigate whether a single contact accelerometer device could sufficiently measure dynamic changes of the sympathetic nervous system (PEP) and preload effects on the heart (LVET) as compared to the 8 electrode ICG. Overall, the results indicate a positive relationship between the ACC and ICG’s values of PEP and LVET. This study demonstrates that the simpler ACC could act as a complementary method of quantifying event-related changes in both PEP and LVET, therefore introducing a protocol that is sensitive to dynamic underlying physiological processes, accessible to more researchers, and takes about a quarter of the time compared to ICG procedures and analysis.

Notably, the exact moment of opening and closing of the aortic valve within the raw ACC waveform was not readily decipherable prior to the application of appropriate signal filters (Figure 3). While two separate pulses are clearly visible (the first for the opening of the aortic valve, and the second for the closing), it is not as apparent which peak/trough to choose when analyzing the data. To solve this, we applied the filters described in the methods (detailed under “ACC Signal Filtering”) post-collection to create two distinct and smooth peaks to place the representative points in analysis. These filters were consistent and reliable across all participants and trials. Following this filtering, the automated analysis pipeline, MEAP, was able to consistently label the opening and closing of the valves with little to no outliers. While these filters are likely to introduce a bit of delay, we found this to be acceptable considering that the delay would be a fixed offset within participants. The delay effect should be considered when these filters are applied to data in experiments that are not solely concerned with relative changes of autonomic activity within individuals. If other researchers consider running these filters in real-time, they should be mindful of an added delay, particularly from the FIR filter. If researchers are concerned about the alterations that an FIR filter might make to the PEP and LVET values, alternative filtering methods of choice can be readily applied.

While PEP measured by ICG during the baseline could be predicted from the ACC with a linear mixed effect model (p<0.001), the complementary Bayesian model resulted in only marginal significance. This suggests that a subset of participants might be driving the results in the linear mixed effects model. It is important to recognize that in the baseline resting condition the natural variation of PEP is very low. Furthermore there may be sources of noise that are different in the two measures. This suggests that the ACC measure might not be suitable for measuring minor fluctuations of baseline PEP within a given subject. Alternatively, the difference between ACC and ICG measures of PEP at rest may also diminish with a larger sample size. To help account for the limited sample size, Bayesian models provided the addition of a more informative prior for studies that try to replicate or build upon our effect. Future studies will need to test the robustness of our observed effects with a larger sample size.

In this study, we tested whether the ACC device is capable of acting as a reliable complement to the ICG’s measure of the mechanistic movement of the aortic valve. While calculating PEP required indices from two apparatuses (ICG and ECG), LVET is quantified as the time period between the opening and closing of the aortic valve (requiring only ICG). LVET therefore provides a more reliable comparison between ACC and ICG. Importantly, our results indicate a strong positive relationship between the ACC and ICG’s LVET measures for both baseline and Valsalva trials, suggesting that this device may be used to measure both minor and major fluctuations in preload effects on the heart.

It should be noted that the ACC and ICG signals arise from different mechanisms, resulting in differing lengths of PEP and LVET per method, as observed in the tables 1 and 2. While the ICG signal is an impedance measure of electrical activity driven by change in the orientation of red blood cells with blood flow velocity, the ACC is a mechanical effect of a pressure wave that the heart is generating. Light travels faster than sound, therefore the electrical signal from ICG is faster than the mechanical signal resulting from the ACC. This bias remains constant across both baseline and Valsalva conditions, suggesting that these differences are resulting from slightly different physiological measures. Because of this difference, experimenters should put this under consideration when comparing raw PEP and LVET values resulting from ACC to the more typical ICG raw PEP and LVET. We recommend the use of the accelerometer for measuring a relative change in PEP and LVET over time in response to physiologic event changes, such as in detecting changes of sympathetic drive and ANS stress response.

An alternative approach to derive measures of aortic movement is with a phonocardiogram. In comparison to a phonocardiogram, which measures sound waves related to valve closure or blood flow, the ACC isolates the acceleration of underlying tissue along three spatial axis (X, Y, Z) simultaneously. By choosing the appropriate axes, cardiac valve opening and closure activity can be reliably identified. An advantage of using acceleration is that it has a flat frequency response at low frequencies whereas a phonocardiogram does not. The ACC is more sensitive at detecting low frequency components associated with valve closure. Furthermore, phonocardiograms come in many different masses, sizes and frequency responses, and their “in-band” frequency response depends greatly on the coupling (attachment) strategies from the phonocardiogram to the skin’s surface. Meanwhile, the ACC technology can be configured to be highly repeatable over a range of ACC chips, assuming the chips have similar acceleration ranges and signal-to-noise ratios (Durand & Pibarot, 1995).

The wireless nature of the ACC device discussed in the present study provides the potential to be used to measure ANS changes in ambulatory tasks, such as with the use of a virtual reality system. For ambulatory or other active performance tasks, an ECG configuration with the red ECG at a more stable body location (e.g. rib cage) as opposed to the triceps, may produce results that are more resilient to movement artifact. The use of other ECG placements are to the discretion of the investigator, as this ACC method should be stable across any reliable ECG configuration. Future experimentation is required to determine the reliability of this application in this manner. Additionally, the SNS response is subject to a predictive “top-down” cognitive control, making it especially interesting to examine simultaneously with fMRI activity (Critchley, 2005; Shoemaker et al., 2012). The combination of impedance cardiography with fMRI has recently been introduced and is gaining increased interest, yet such a study is yet to be done with the use of an accelerometer (Cieslak et al., 2015). Our results suggest that the ACC works reliably in the supine position that is required inside the MRI, therefore this should be feasible once an MRI compatible system is validated. We encourage further research to compare these ACC methods during psychological and other physical test conditions that are known to stimulate ANS and more specifically, SNS, effects on the heart.

In conclusion, we saw a clear coordination between measures of the SNS and preload effects on the heart using classic impedance cardiography and our ACC in response to a reliable physiological perturbation. As a result, we believe the ACC device introduces a simple procedure to measure changes of the ANS, with easily analyzable output and the potential to be used at a greater capacity than current impedance cardiography methods for measuring these changes.

## ACKNOWLEDGEMENTS

The research was supported by award #W911NF-16-1-0474 from the Army Research Office and by the Institute for Collaborative Biotechnologies under Cooperative Agreement W911NF-19-2-0026 with the Army Research Office.

## IMPACT STATEMENT

Our research presents a wireless accelerometer (ACC) device as a complementary method to measure pre-ejection period (PEP) and left ventricular ejection time (LVET) to quantify changes of the autonomic nervous system. Obvious and significant associations were found between impedance cardiography (ICG) and ACC estimates of PEP and LVET as participants engaged in a classic physical stress task. As compared to current methods of ICG, the ACC cuts down on electrode preparation and data analysis time and is accessible to more researchers.

