## Supplementary Materials for "An Accelerometer Based Heart Monitor to Measure Changes of the Autonomic Nervous System"

*Power coherence analysis to compare respiration signals between the ACC and ICG's Z0.*

A relative benefit of ICG is the ability to retrieve respiration from the raw impedance (Z0) channel. While our study was not designed to evaluate respiratory signals with (ACC), in response to a reviewer query we tested whether respiration information could in fact be derived from the accelerometer by using impedance cardiograph (ICG) rather than direct respiration measurements as the reference. We calculated a nonparametric regression (Spearman's rho) between the signal envelopes of the two band-passed signals. These signal envelopes were computed on the bandpassed signals (0.1-0.5Hz) using the root mean squared values in sliding windows of 1000 samples (10s of data). Testing the distribution of rhos (i.e., across subjects) against 0, separately for each condition, we see agreement in the Valsalva condition ($p < 0.001$, 95% CI [0.231, 0.469]), and a trend towards agreement within both baseline ($p = 0.21$, 95% CI [-0.069, 0.292]) and supine ($p = 0.135$, 95% CI [-0.051, 0.329]) conditions. Similar to our findings with PEP and LVET, these weaker agreements in baseline and supine conditions may diminish with a larger sample size.

*Inter-autonomic Criterion Validity*

Inspired by a reviewer's suggestion, we've included comparisons between heart rate variability (HRV) and PEP and LVET derived from both the ACC and ICG during the Valsalva condition. We use broadband HRV (the variance in sliding 20 second windows) across the whole session, to provide one estimate per second in time for each subject. We then perform the same procedure as we did for estimating criterion validity (see manuscript), but instead predicting HRV separately from both ICG and ACC. We observe fairly robust effects driven by the Valsalva from the start of the maneuver at 60s. The HRV response is credibly anti-correlated with PEP, estimated with both the ICG and ACC sensors (as observed by the agreement histograms). HRV associations are also negatively correlated with LVET. The Valsalva maneuver triggers a large variety of physiological changes in the body, resulting in two different physiological effects (the Valsalva and its recovery). Due to this, we refrain from making assumptions of how HF HRV should relate to the PEP and LVET during this procedure, but rather use this analysis to point out the strong correlations between these measures.

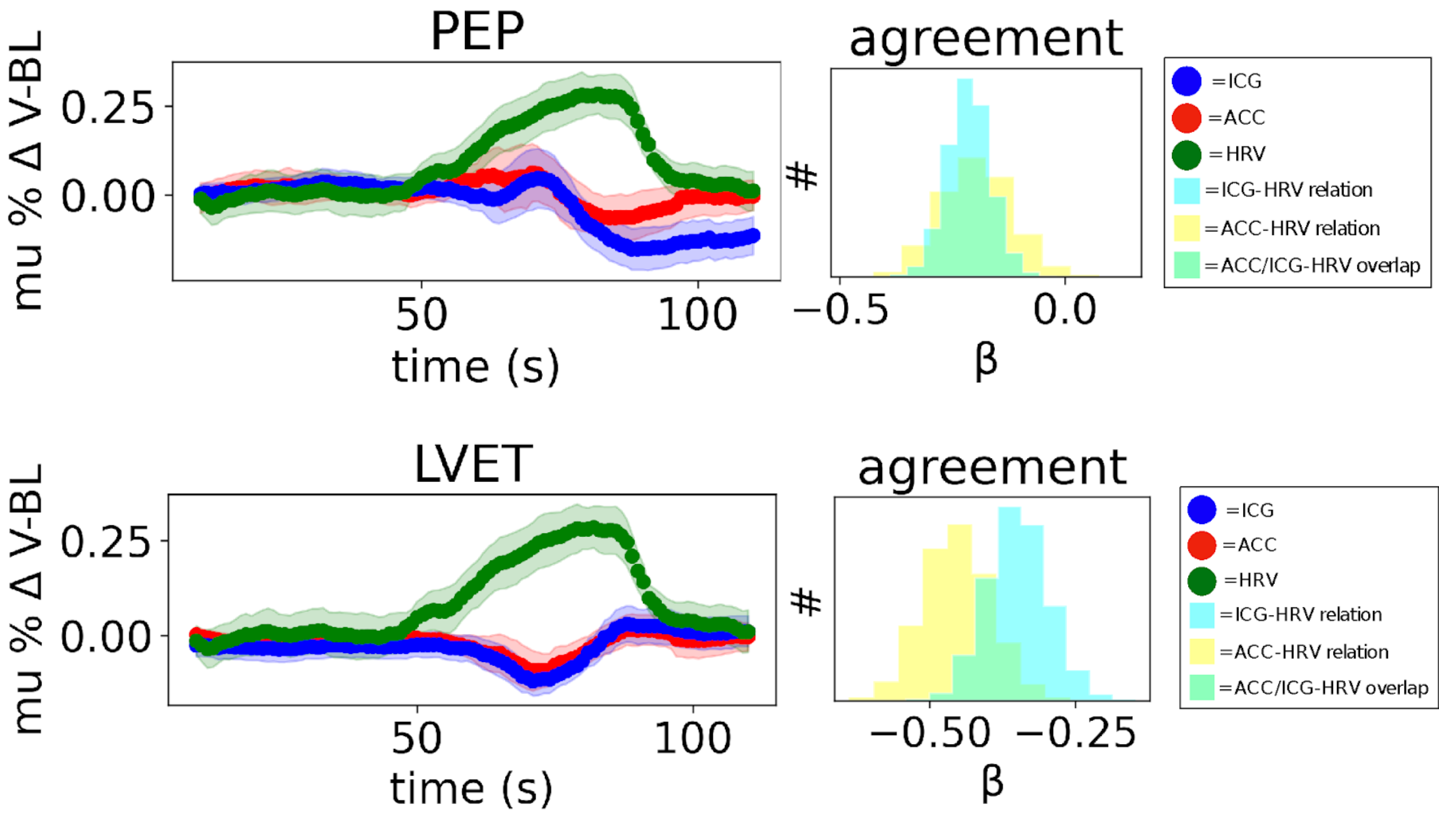


*Internal Consistency Pairwise Departures*

In addition to our internal consistency plots in our manuscript, we perform an exhaustive departure analysis (matrix) to test whether variability at any given moment is credibly different to that at any other moment (non-overlapping HDIs) for each of the eight measurements - two conditions (BL,V), two sensors (ACC, ICG) and two physiology measures (PEP, LVET). For ICG (blue) and ACC (red), dots show the mean of the posterior estimating the standard deviation across subjects at each data point, while the shaded region depicts the HDI, i.e., the credible range of variability across subjects at each moment in time. Within each of the eight measurements, we see no strong evidence that the stand deviations across subjects at any timepoint was credibly different to those at all other timepoints.

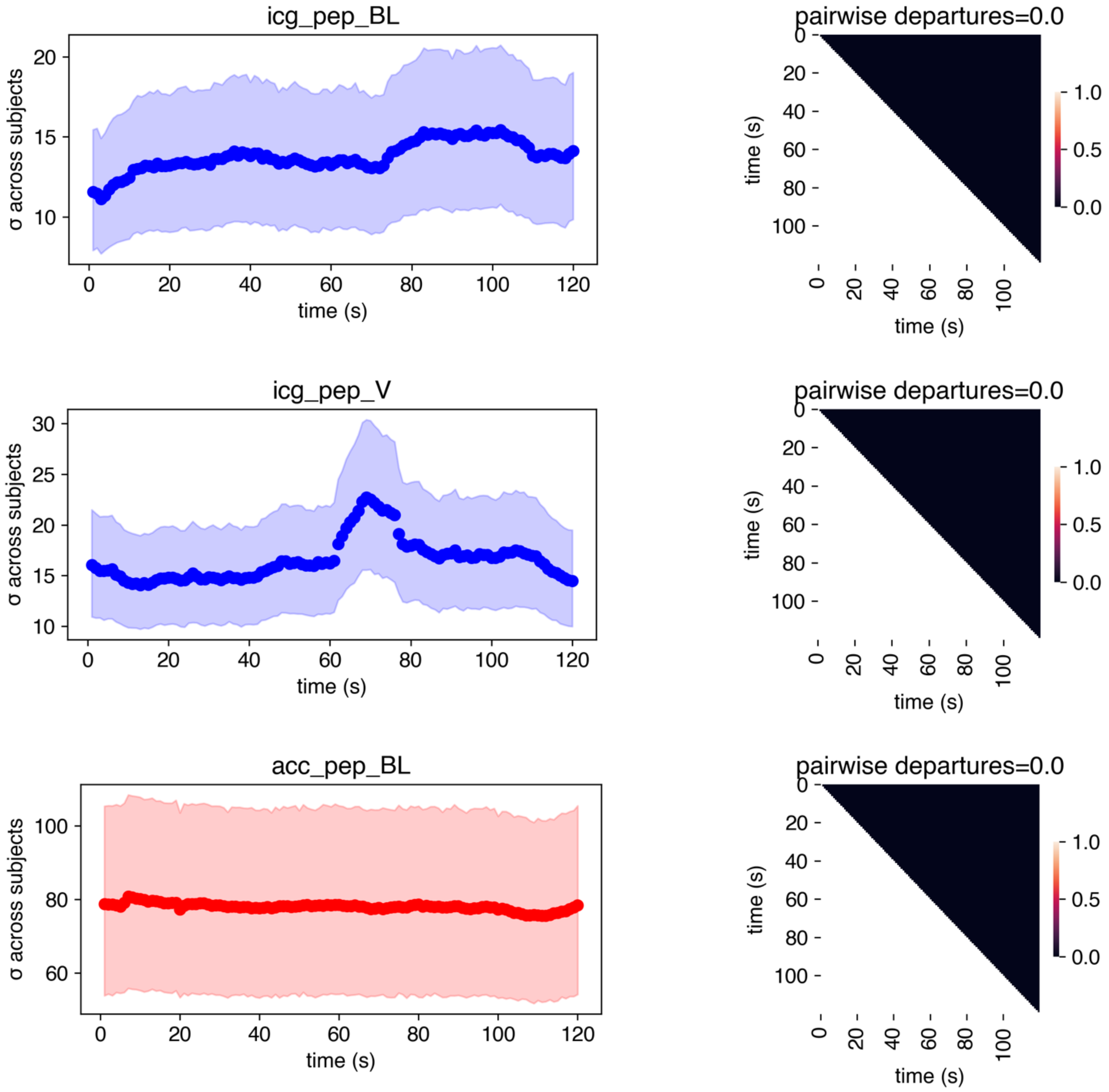

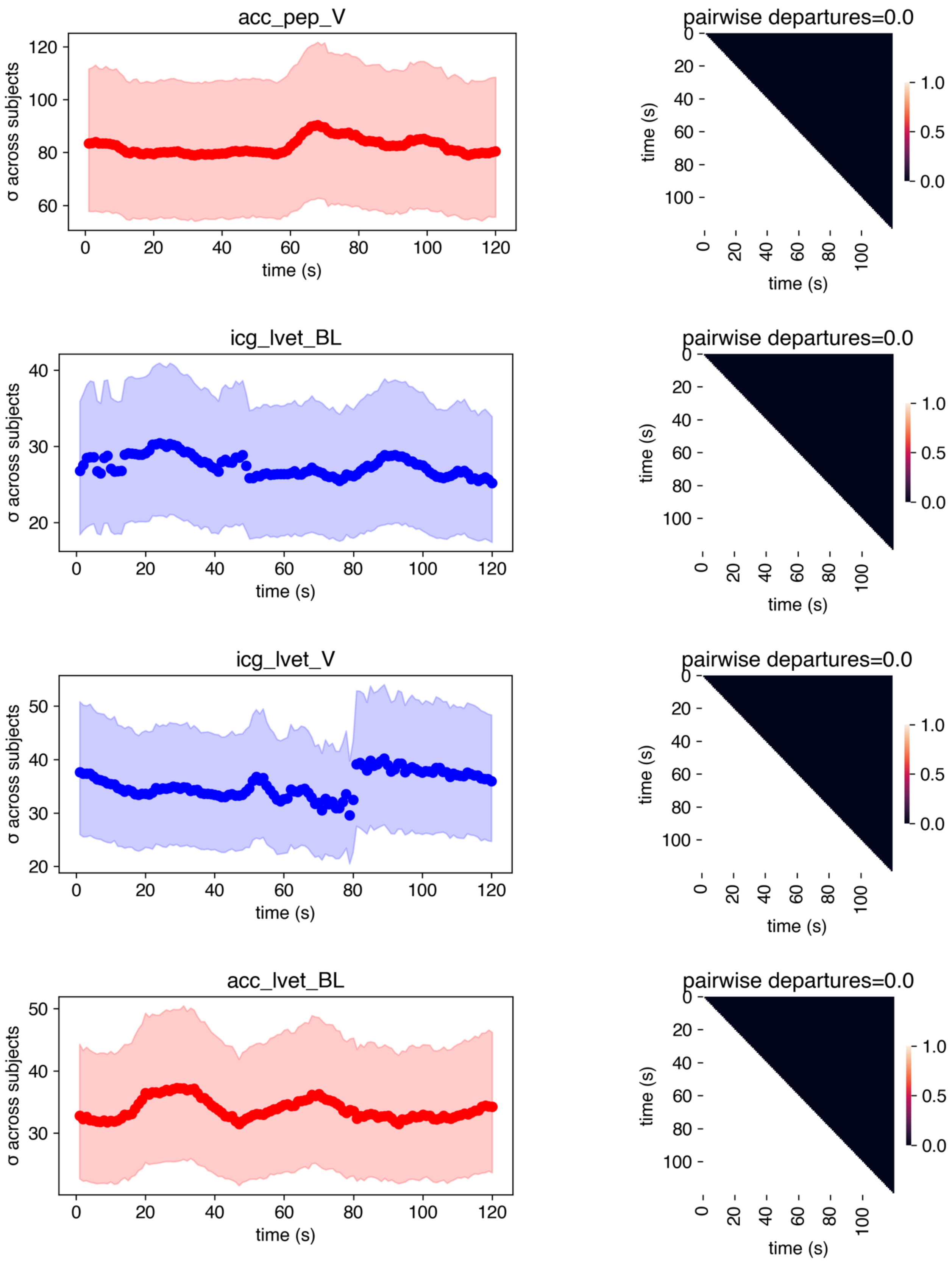
acc_pep_V
σ across subjects
time (s)
pairwise departures=0.0
time (s)
icg_lvet_BL
σ across subjects
time (s)
pairwise departures=0.0
time (s)
icg_lvet_V
σ across subjects
time (s)
pairwise departures=0.0
time (s)
acc_lvet_BL
σ across subjects
time (s)
pairwise departures=0.0
time (s)
1.0
0.5
0.0

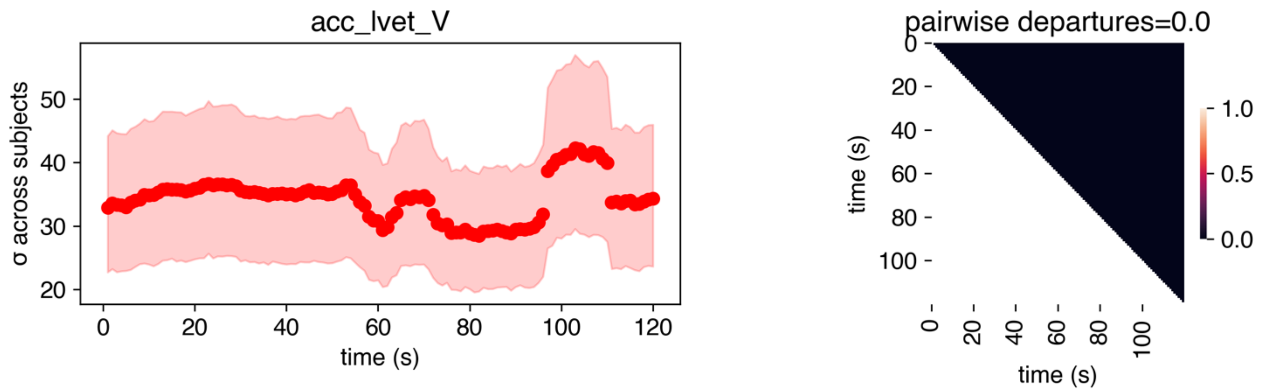


*Bland Altman Plots*

In response to an editor query, we include Bland-Altman plots on the PEP and LVET delta timeseries. These plots depict the point-by-point average of recordings with the ACC and ICG, plotted against their difference, such that skew in the data reflects agreement that depends on the magnitude of recorded values. Alongside each subject's plot is a histogram of delta values, which show normal distributions in most cases.

### Baseline PEP

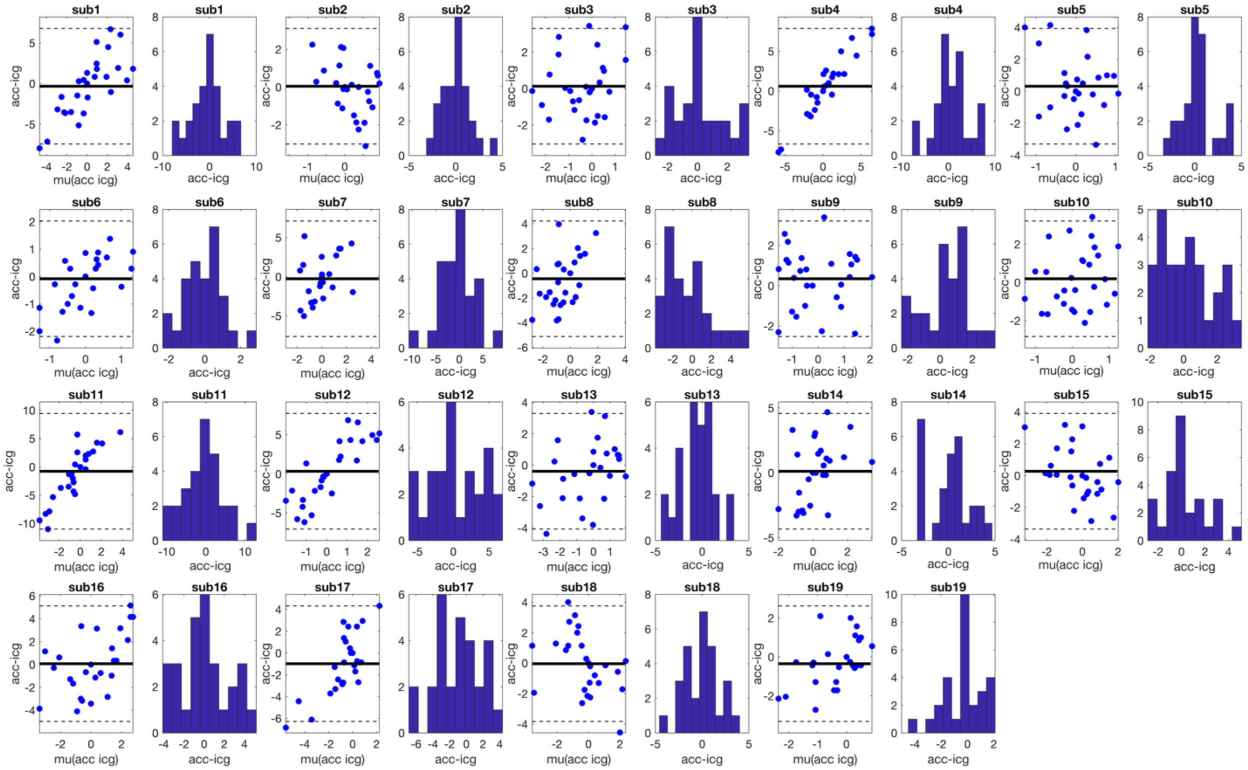

### Supine PEP

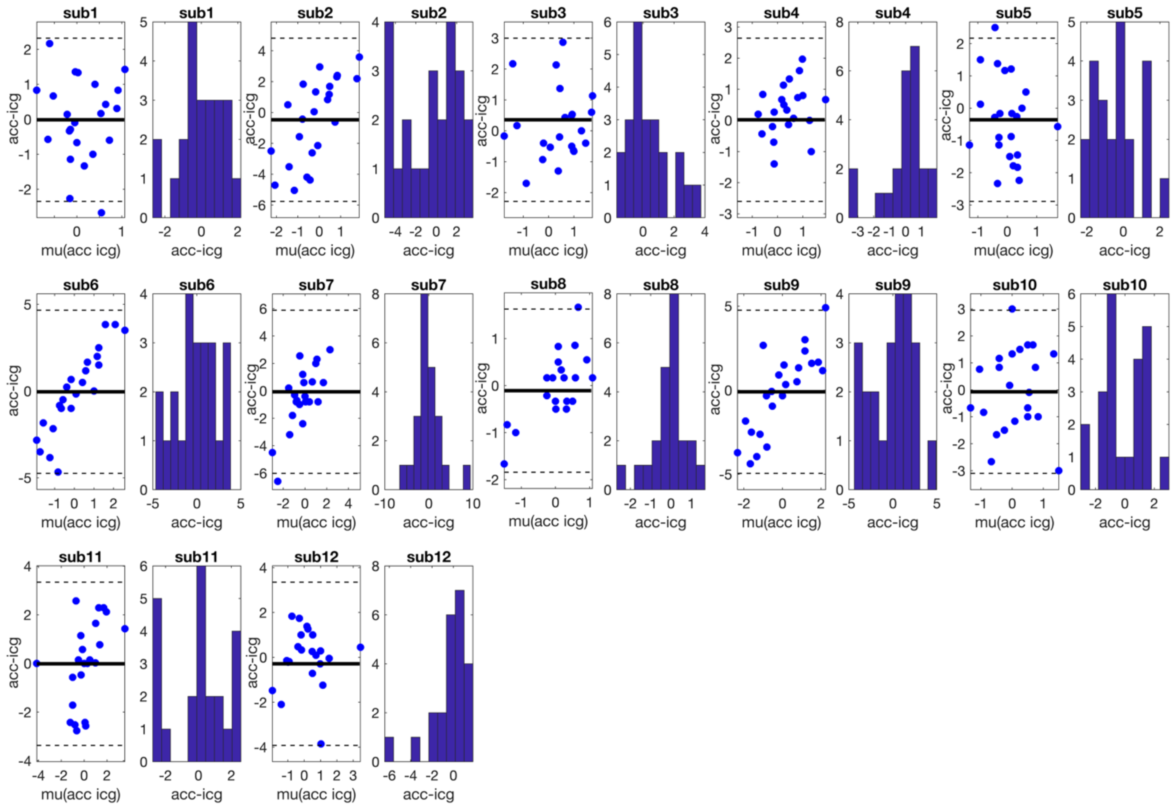
sub1
sub1
sub2
sub2
sub3
sub3
sub4
sub4
sub5
sub5
sub6
sub6
sub7
sub7
sub8
sub8
sub9
sub9
sub10
sub10
sub11
sub11
sub12
sub12
acc-icg
mu(acc icg)

### Valsalva PEP

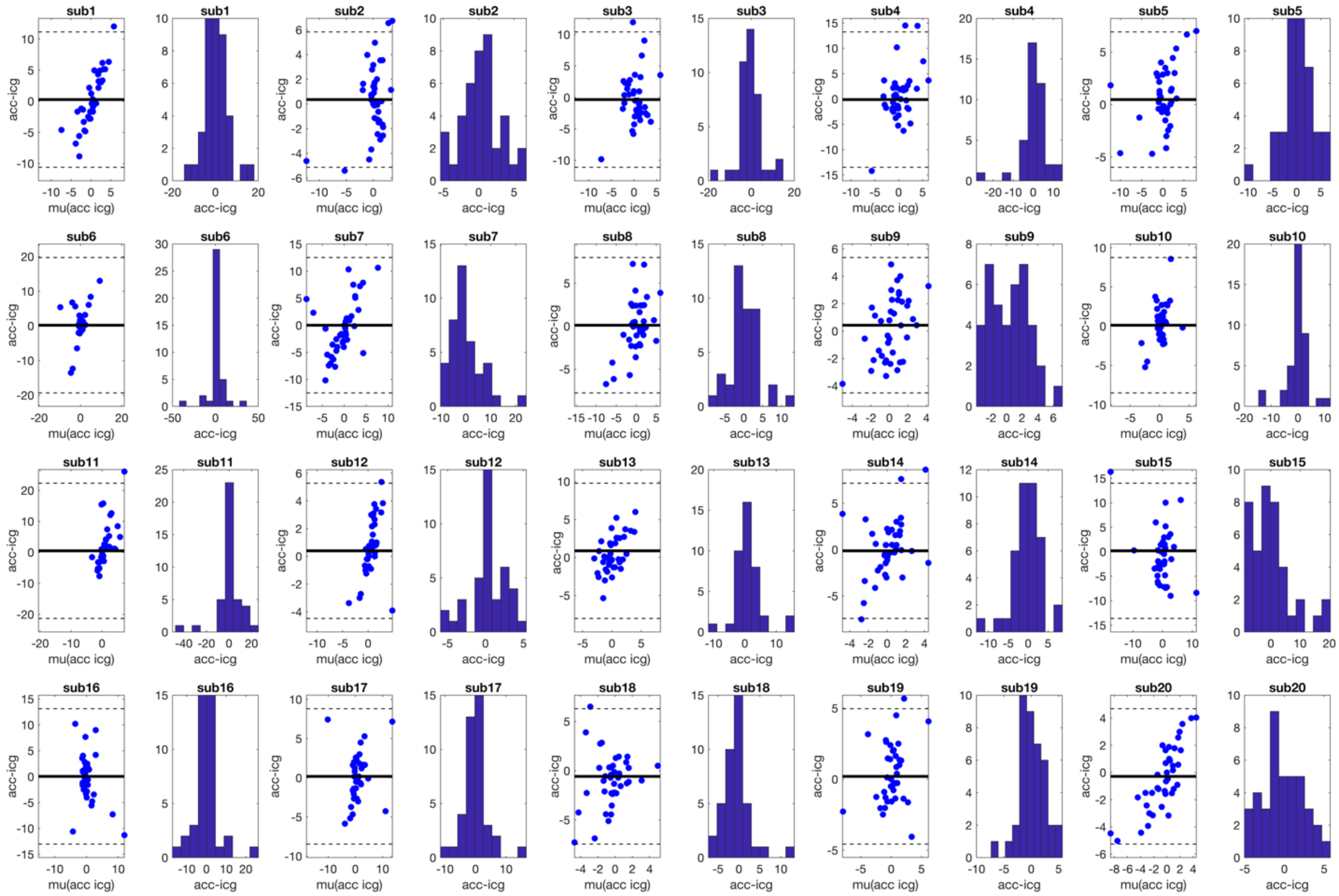

#### Baseline LVET

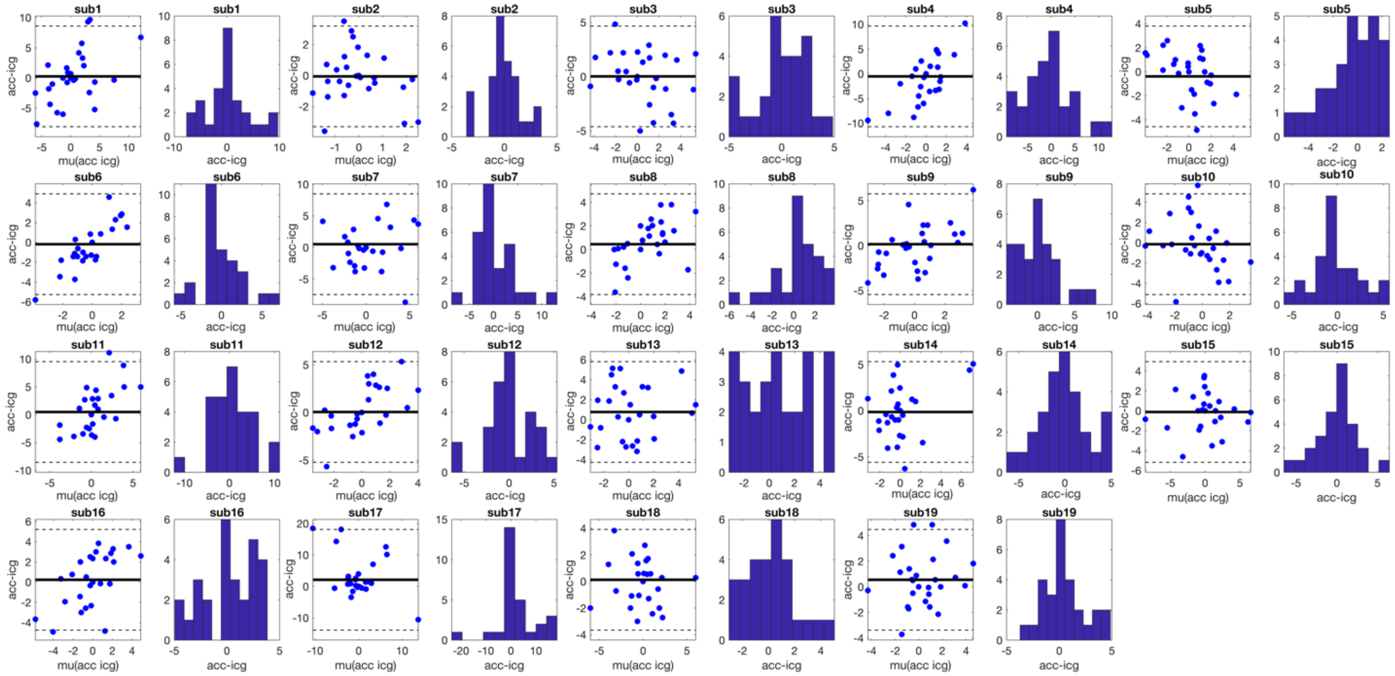


#### Supine LVET

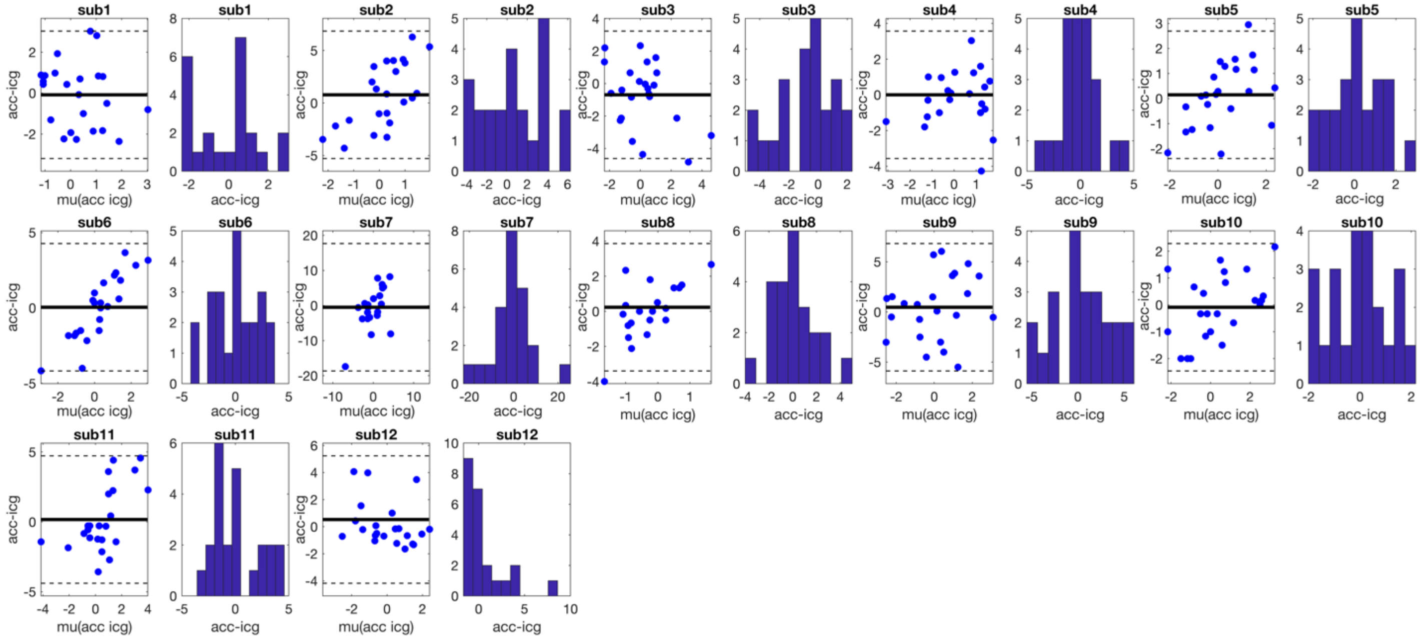

### Valsalva LVET

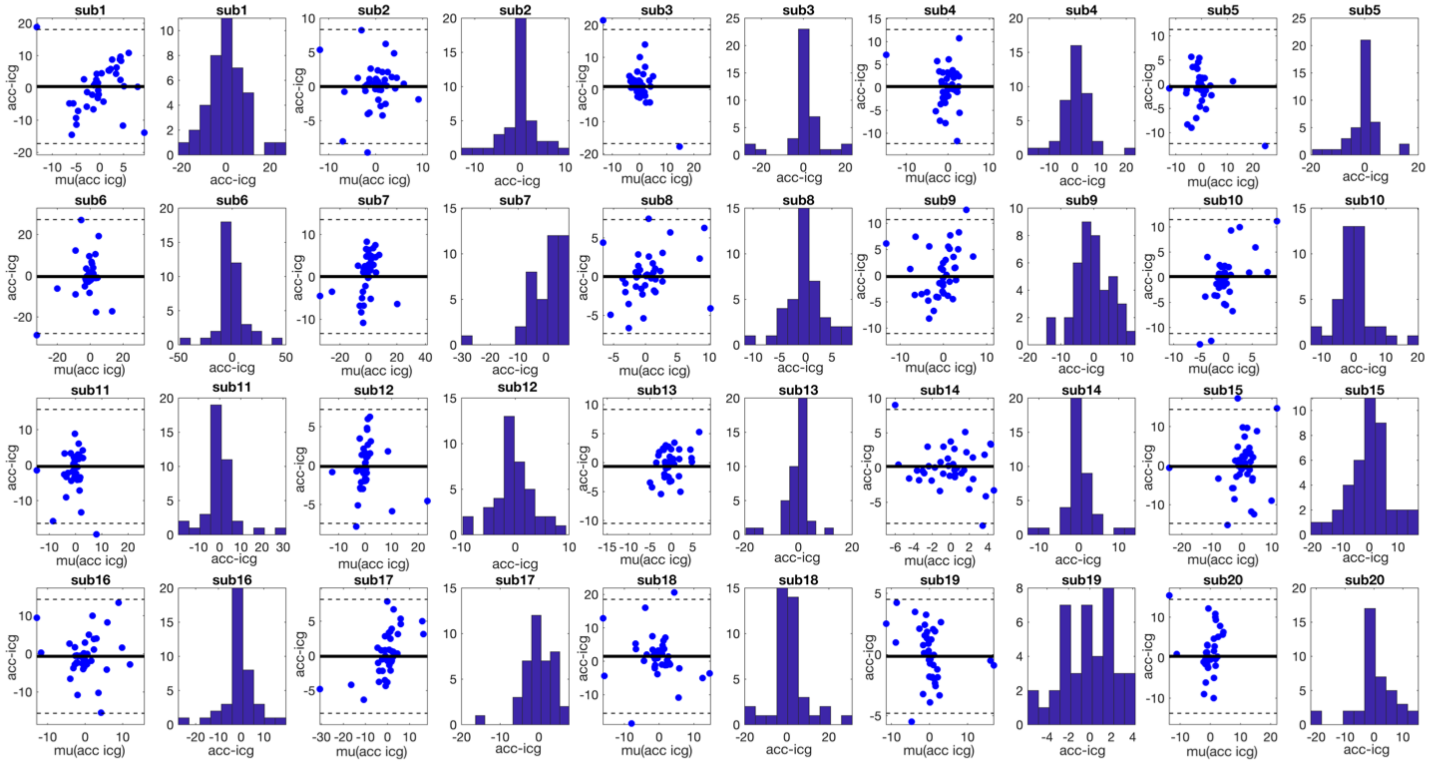
